# Cross-species analysis identifies conserved transcriptional mechanisms of neutrophil maturation

**DOI:** 10.1101/2022.11.28.518146

**Authors:** Stefanie Kirchberger, Mohamed R. Shoeb, Daria Lazic, Kristin Fischer, Lisa E. Shaw, Filomena Nogueira, Fikret Rifatbegovic, Eva Bozsaky, Ruth Ladenstein, Bernd Bodenmiller, Thomas Lion, David Traver, Matthias Farlik, Sabine Taschner-Mandl, Florian Halbritter, Martin Distel

**Author notes:** contributed equally (Stefanie Kirchberger, Mohamed R. Shoeb). equally contributing last authors (Florian Halbritter, Martin Distel). **Corresponding authors:**, F+43 1 40470 7150, **Address:** CCRI, Zimmermannplatz 10, 1090 Vienna, Austria.

## Abstract

Neutrophils are evolutionarily conserved innate defense cells implicated in diverse pathological processes. Zebrafish models have contributed substantially to our understanding of neutrophil functions, but similarities to human neutrophil maturation have not been characterized limiting applicability to study human disease.

We generated transgenic zebrafish strains to distinguish neutrophil maturation grades *in vivo* and established a high-resolution transcriptional profile of neutrophil maturation. We linked gene expression at each stage to characteristic transcription factors, including C/ebpβ, important for late neutrophil maturation. Cross-species comparison of zebrafish, mouse, and human confirmed high molecular similarity in immature stages and discriminated zebrafish-specific from pan-species gene signatures. Applying pan-species neutrophil maturation signatures in RNA-seq data from neuroblastoma patients revealed an association of metastatic tumor cell infiltration in the bone marrow with an increase in mature neutrophils.

Our detailed neutrophil maturation atlas provides a valuable resource for studying neutrophil function at different stages across species in health and disease.

**Graphical abstract:** 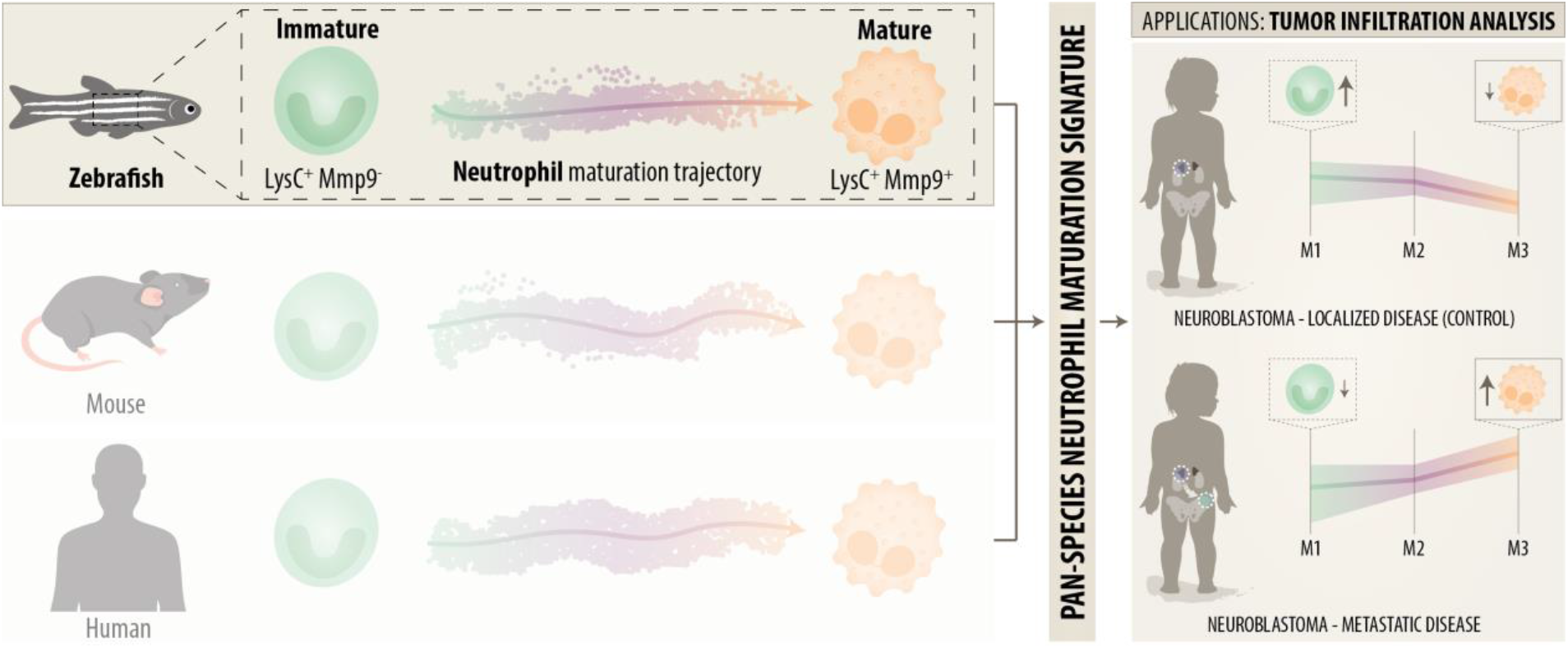

## Introduction

Neutrophils are the most abundant immune cell population in humans and the first responders to injury and infection^1, 2^. In mammals, neutrophils mature in the bone marrow (BM), during the final steps of a cascade where hematopoietic stem cells differentiate through a granulocyte-monocyte progenitor towards neutrophils^3, 4^. Characteristic granules form throughout development from the promyelocyte to segmented neutrophil stage^5, 6, 7^. Transcription factors of the C/EBP family are key in the formation of granule enzymes^8, 9^. Surface marker expression, nuclear morphology, and granule content have long been used to define neutrophil maturation stages, but recent studies using single-cell RNA sequencing (scRNA-seq) and proteome analysis (CyTOF) have questioned this classification scheme instead suggesting a sequence of continuous transcriptional stages^10, 11, 12, 13, 14, 15^.

Functional diversity of neutrophils at different maturation grades is still understudied^16, 17^. A potential role for different maturation states becomes apparent in cancer, where immature neutrophils often accumulate in blood and tumors, and can have altered effector functions such as reduced phagocytosis, ROS production, NETosis, granularity, chemokine receptor expression and increased suppressive functions^6, 16, 18^. Neutrophils are now known to be involved in almost every stage of cancer such as tumor initiation by ROS production, growth, angiogenesis, and the conditioning of the pre-metastatic niche^6, 17^. Different tumor-associated neutrophil (TAN) populations polarized by TGFβ or IFNβ towards pro- or anti-tumor roles have been observed. Interestingly, their opposing roles in cancer progression have been linked to different maturation stages and densities^18, 19, 20^.

Zebrafish are a versatile model to study neutrophil functions in infection and tumorigenesis thanks to the availability of many fluorescent transgenic lines and the prospect of intravital imaging^21, 22, 23^. Hematopoiesis in zebrafish occurs in the kidney marrow, where all major immune cell types known from humans are present and generally considered evolutionarily conserved^24, 25^. However, to date the transcriptional states of zebrafish neutrophils during maturation and associated functions *in vivo* have not been mapped to their human equivalents, thus limiting cross-species comparisons. Comparative transcriptomic studies have been hampered by under-representation of neutrophils in many human datasets due to technical difficulties and by a lack of suitable zebrafish lines allowing effective sorting of neutrophils comprising multiple maturation stages^26^.

Here, we established *Tg(lysC:CFPNTR)*^*vi002*^/*Tg(BACmmp9:Citrine-CAAX)*^*vi003*^ double transgenic zebrafish, which allowed us to distinguish immature from mature neutrophils *in vivo*, visualize their interactions with bacteria and tumor cells, and isolate maturing neutrophil populations for morphological and transcriptional analysis. We used scRNA-seq to catalogue transcriptional changes during maturation and to identify critical transcription factors, including C/ebpβ, which had previously been thought to be involved only in emergency granulopoiesis^27, 28^. Finally, cross-species comparison enabled the definition of a conserved gene signature, which we applied to analyze bulk human tumor transcriptome data and correlate neutrophil maturation stage with BM metastasis.

## Results

### Mmp9 transgene identifies mature neutrophils in zebrafish

In order to distinguish immature from mature neutrophils *in vivo*, we generated transgenic zebrafish expressing membrane-directed Citrine under the control of regulatory elements for mmp9 (*Tg(BACmmp9:Citrine-CAAX)*^*vi003*^), a tertiary granule protein in mammalian mature neutrophils and expressed in zebrafish mature heterophils^29, 30^. *Tg(BACmmp9:Citrine-CAAX)*^*vi003*^ zebrafish embryos/larvae report *mmp9* transcription from 2 days post fertilization (dpf) in the epithelia of the tail fin and the distal gut **(Supplementary Fig. 1A)**. Additionally, a dotted pattern became apparent along the head, yolk, and in the caudal hematopoietic tissue suggesting expression in a leukocyte population. We confirmed that Citrine fluorescence specifically reports *mmp9* expression by detecting *mmp9* RNA in FACS-purified Citrine^+^ but not in Citrine^-^ cells **(Supplementary Fig. 1B)**.

**Fig. 1:**
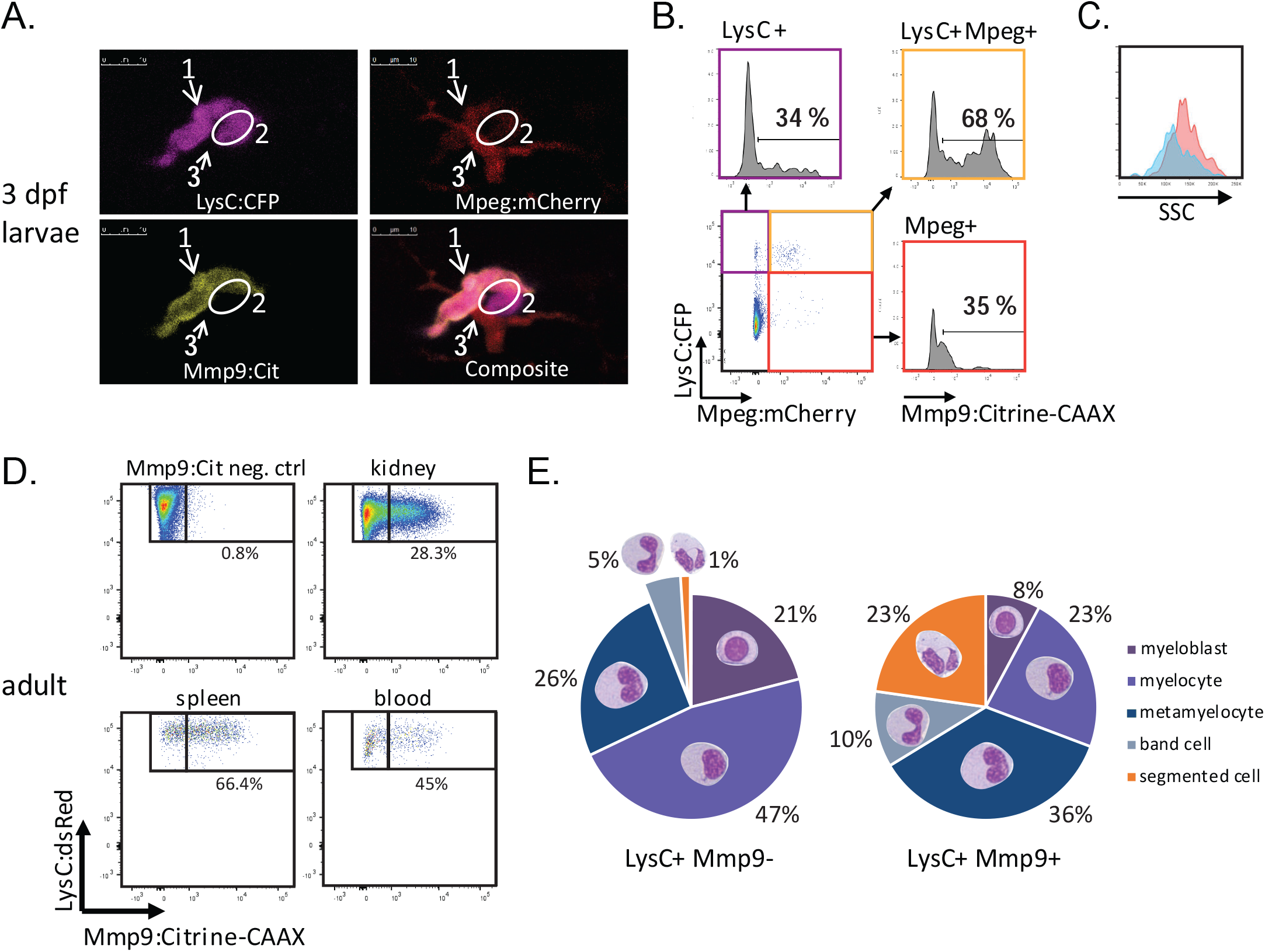
mmp9:Citrine identifies mature neutrophils. **(A)** Confocal images of triple transgenic larvae *Tg(lysC:CFP-NTR)*^*vi002*^ *Tg(BACmmp9:Citrine-CAAX)*^*vi003*^ *Tg(mpeg:mCherry)*^*gl23*^ at 3 dpf reveal myeloid cells with different expression levels of the three analyzed markers. *#1 neutrophil: lysC:CFP*^*+*^ mpeg:mCherry^low^, *mmp9:Citrine*^*+*^; *#2 neutrophil: lysC:CFP*^*+*^ *mpeg:mcherry*^*-*^, *mmp9:Citrine*^*-*^; *#3 macrophage lysC:CFP*^*-*^ *mpeg:mcherry*^*+*^, *mmp9:Citrine*^*-*^. Scale bars 10 µm. Images acquired with a Leica TCS SP8 WLL microscope (HC PL APO CS2 40x/1.10 WATER objective). Maximum projections were performed in the Leica LAS X software. **(B)** Flow cytometric analysis (BD Fortessa) of myeloid cells isolated from a pool of 20 triple transgenic larvae *Tg(lysC:CFP-NTR)*^*vi002*^ *Tg(BACmmp9:Citrine-CAAX)*^*vi003*^ *Tg(mpeg:mCherry)*^*gl23*^ at 3 dpf shows that Mmp9: Citrine predominates in the *lysC:CFP*^*+*^ *mpeg:mcherry*^*low*^ neutrophil population. Dotblot (bottom left) shows the quadrants further analyzed in the histograms for mmp9 expression. n = 2. **(C)** *Mmp9:Citrine*^*+*^ neutrophils (red histogram) have a higher side scatter and therefore granularity compared to *mmp9:Citrine*^*-*^ neutrophils (blue histogram). Flow cytometric analysis of cells isolated from *Tg(lysC:CFP-NTR)*^*vi002*^ *Tg(BACmmp9:Citrine-CAAX)*^*vi003*^ at 2 dpf. Cells were gated on *the lysC:CFP*^*+*^ *mmp9:Citrine*^*+*^ or *lysC:CFP*^*+*^ *mmp9:Citrine*^*-*^ populations and analyzed for SSC. n = 3. **(D)** Cells were isolated from kidneys, spleen or blood of adult *Tg(lysC:dsRed)*^*nz50Tg*^ *Tg(BACmmp9:Citrine-CAAX)*^*vi003*^ and analyzed by flow cytometry for frequency of Mmp9^+^ cells. **(E)** Neutrophils were isolated from kidneys of adult *Tg(lysC:dsRed)*^*nz50Tg*^ *x Tg(BACmmp9:Citrine-CAAX)*^*vi003*^ and sorted into *lysC:dsRed*^*+*^ *mmp9:Citrine*^*+*^ *or lysC:dsRed*^*+*^ *mmp9:Citrine*^*-*^ populations, cytospins were prepared and Pappenheim stained. Different neutrophil maturation stages were scored blinded, showing that the *lysC:dsRed*^*+*^ *mmp9:Citrine*^*+*^ population consists of highly differentiated neutrophil stages. Graph presents average percentages of n = 4 kidneys, 721 Mmp9^+^ cells and 845 Mmp9^-^ cells were analyzed in total.

To examine the identity of Mmp9^+^ leukocytes, mmp9:Citrine fish were crossed with fish double-transgenic for myeloid markers *Tg(lysC:CFP-NTR)*^*vi002*^ (labelling neutrophils), and *Tg(mpeg:mCherry)*^*gl23*^ (macrophages) ^31^. Live imaging of triple-transgenic larvae at 3 dpf revealed the existence of Mmp9^-^ and Mmp9^+^ subpopulations of lysC:CFP^+^ neutrophils **(Fig. 1A)**. Mmp9^+^ cells had only low mpeg:mCherry expression, while *bona-fide* macrophages were highly positive for mpeg:mCherry and showed no detectable Mmp9 expression. Complementary analysis by flow cytometry confirmed the highest frequency of Mmp9-expressing cells in the LysC^+^Mpeg^lo^ neutrophil (68%, Q2) population **(Fig. 1B)**. Only few Citrine-positive cells were detected in LysC^+^Mpeg^-^ neutrophils (34%, Q1) and Mpeg^hi^LysC^-^ macrophages (35%, Q3). Mmp9^+^ cells showed an increased side scatter (SSC) compared to Mmp9^-^ cells indicating a higher granularity and intracellular complexity, which suggests a more mature phenotype **(Fig. 1C)**.

Time-lapse imaging of cellular behavior after wounding showed that both, LysC^+^Mmp9^+^ and LysC^+^Mmp9^-^ neutrophils had a similar round morphology and moved to and from the wound quickly with amoeboid motility as described for neutrophils^32^ **(Movie 1)**. In contrast, Mpeg^+^ cells showed the protrusions typical of macrophages and stayed close to the wound edge.

In adult zebrafish LysC^+^Mmp9^+^neutrophils were detectable in the whole kidney marrow (WKM), the primary hematopoietic organ of teleost fish, as well as in the spleen and blood **(Fig. 1D)**. We investigated morphology and maturation grade of sorted LysC^+^Mmp9^+^and LysC^+^Mmp9^-^ populations from WKM on stained cytospins **(Fig. 1E)**. Only 1% of Mmp9^-^ cells contained segmented nuclei compared to 23% of the Mmp9^+^ fraction, indicating an advanced maturation grade of the latter. Collectively, our data show that Mmp9 is a suitable marker for mature neutrophils in zebrafish consistent with previous data for human cells^33^.

### Mmp9+ neutrophils are highly phagocytic and rapidly recruited to wounds

The phagocytic capacity of neutrophils increases with their maturation grade^34^. We performed *in vivo* phagocytosis assays by injecting live mCherry-labelled *E. coli* into 2 dpf *Tg(lysC:CFPNTR)*^*vi002*^*/ Tg(BACmmp9:Citrine-CAAX)*^*vi003*^ larvae. Both neutrophil subpopulations, LysC^+^Mmp9^-^ and LysC^+^ Mmp9^+^, were able to ingest bacteria **(Fig. 2A)**. Quantification of *E. coli* uptake as seen by overlap of mCherry (*E. coli*) and CFP (neutrophil) fluorescence by flow cytometry showed a significantly higher percentage of Mmp9^+^ neutrophils with bacterial cargo compared to Mmp9^-^ neutrophils (mean= 23.8% and 10.7%, respectively; n = 5; paired t-test, *P= 0*.*007*) **(Fig. 2B, C)**. Furthermore, Mmp9^+^ neutrophils were more efficiently recruited to sites of bacterial infection (67.3% vs. 32.7%; n = 33; paired t-test; *P = 0*.*002*) **(Fig. 2D)** and the overall frequency of Mmp9^+^ cells was increased during infection (mean= 56.4% and 39.2% in infected and PBS, respectively; n = 5; paired t-test; *P = 0*.*009*) **(Supplementary Fig. 2A, B)**.

**Fig. 2:**
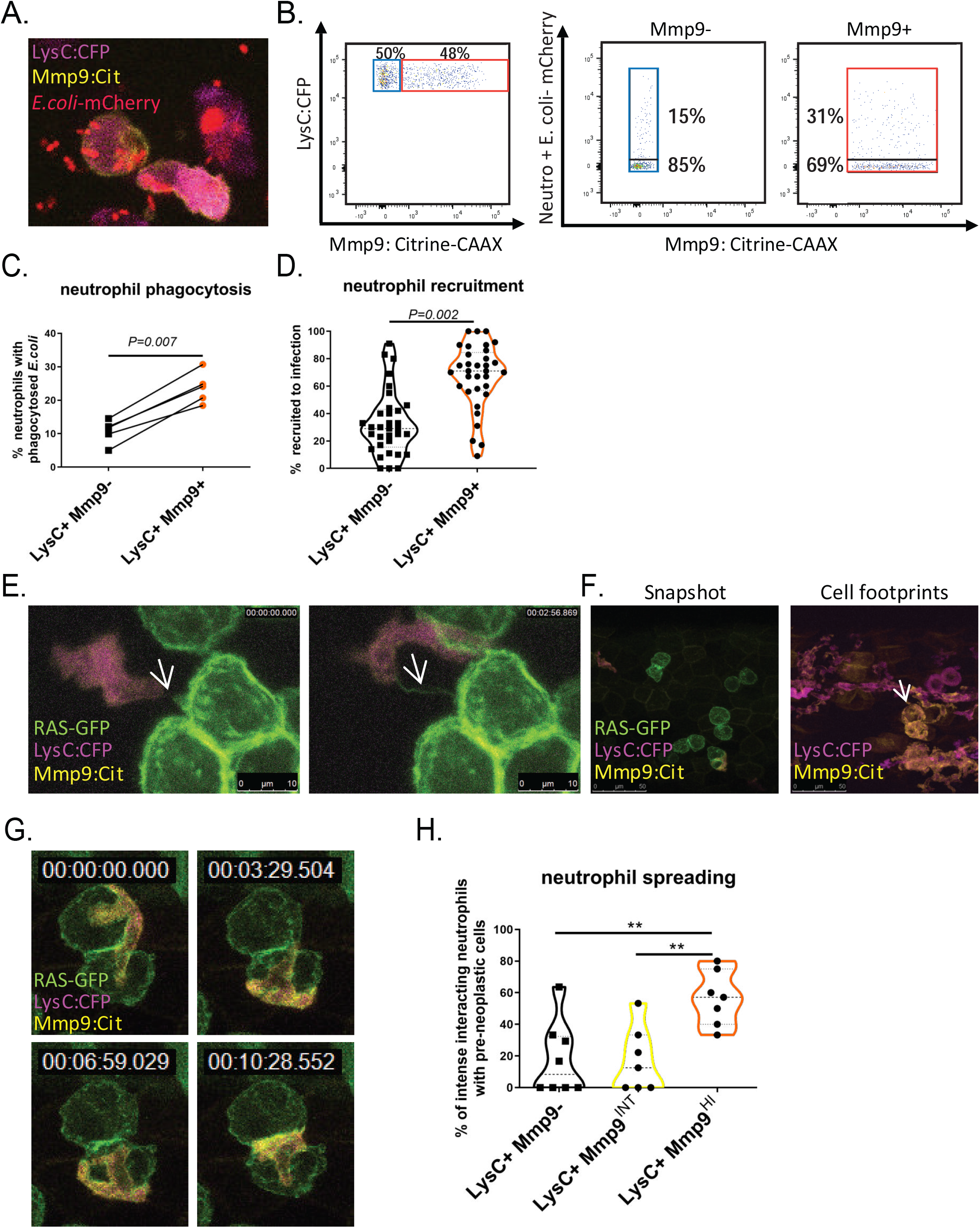
Mmp9+ neutrophils show functions of mature neutrophils. Images were acquired with a HC PL APO CS2 40x/1.10 WATER objective on a Leica Sp8 confocal microscope using LAS X software. **(A-C)** *In vivo* phagocytosis assays were performed by injecting mCherry-labelled *E. coli* into the caudal vein of *Tg(lysC:CFP-NTR)*^*vi002*^ *Tg(BACmmp9:Citrine-CAAX)*^*vi003*^ zebrafish larvae at 2 dpf. **(A)** *E. coli* were observed inside both, *lysC:CFP*^*+*^ *mmp9:Citrine*^*+*^ *and lysC:CFP*^*+*^ *mmp9:Citrine*^*-*^ neutrophils 6 hpi. **(B)** Representative flow cytometry plots of cells isolated from *E. coli* injected larvae 6 hpi, pre-gated on *lysC:CFP*^*+*^ cells (left) and further gated on Mmp9^-^ (middle) and Mmp9^+^ (right) populations, showing a higher percentage of Mmp9^+^ cells with bacterial cargo. **(C)** Graph summarizing results of five independent phagocytosis experiments analyzed by flow cytometry with pools of approx. 20 larvae each group. Paired t-test was performed; **(D)** neutrophil recruitment to *E*.*coli*-mCherry injected into the otic vesicle of 3 dpf larvae was analyzed by confocal microscopy. **(E-H)** Neutrophil-pre-neoplastic cell interactions were observed by confocal microscopy in *Et(kita:GAL4)*^*hzm1*^ *Tg(UAS:EGFP-HRAS_G12V)*^*io006*^ *Tg(lysC:CFP-NTR)*^*vi002*^ *Tg(BACmmp9:Citrine-CAAX)*^*vi003*^ zebrafish larvae starting at 78 hpf. **(E)** Still Images of z-stack maximum projections from a time-lapse movie showing Mmp9+ neutrophils forming dynamic contacts with GFP+ kita tumor cells. White arrows point at GFP+ cell tether. Z-stacks were acquired every 88 s. Scale bar = 10 µm. **(F)** Snapshots and cell footprints were taken from the same time-lapse movie (5 h movie-time). Superimposition of *lysC:CFP* and *mmp9:Citrine* from all time frames generating neutrophil footprints (right). White arrow points out how the movements of an Mmp9^+^ neutrophil copied the outline of the RAS-GFP+ cluster seen in the snapshot (left). **(G)** Close-up clippings showing an Mmp9^+^ neutrophil spreading over the tumor cell cluster marked by a white arrow in (F). **(H)** Quantification of neutrophil-tumor interactions were performed from maximum projections of different time-lapse movies (n = 8). The frequency of interacting neutrophils of each subpopulation (no, intermediate or high *mmp9:Citrine* levels) getting into close, intense interactions with GFP+ kita/RAS skin pre-neoplastic cells. One-way ANOVA was performed. **p<0.01

Next, we examined neutrophils in a model of pre-neoplastic melanoma (*Et(kita:GAL4)*^*hzm1*^*x Tg(HRAS_G12V:UAS:CFP)*^*vi004*^)^35^. As during infection, we found an increase in Mmp9^+^ cells (mean 46.0% versus 37.4%; in the presence of HRAS^G12V^ versus controls; n = 5; paired t-test; *P = 0*.*046*) **(Supplementary Fig. 2C)**. Live imaging in transparent zebrafish larvae showed that some Mmp9^+^ cells stayed in contact with transformed cells by thin tethers **(Fig. 2E)** and others interacted over a long period **(Movie 2)**, consistent with previous data for LysC^+^ neutrophils^36^. We found that some neutrophils got into close contact, spread out, and crawled over HRAS^G12V+^ cells, seemingly scanning their surface **(Fig. 2G)**. Cell footprints of those neutrophil movements trace the outline of HRAS^G12V+^ clusters **(Fig. 2F)**. Notably, this spreading behavior around HRAS^G12V+^ cells was significantly enriched in Mmp9^HI^ cells (mean 56.5%; n = 8; one-way ANOVA, *P = 0*.*0022* compared to Mmp9^INT^ (17.3%) or Mmp9^-^ (17.9%) cells **(Fig. 2H, Movie 3)**.

In summary, we observed that *mmp9* transgene expression correlates with a high maturation grade in neutrophils with increased effector functions, such as recruitment and phagocytosis during bacterial infections and augmented interactions with pre-neoplastic cells.

### Single-cell transcriptomics defines zebrafish neutrophil maturation stages

To scrutinize neutrophil maturation in zebrafish at the molecular level and to enable cross-species comparison to mammals, we performed droplet-based scRNA-seq on WKM and on LysC^+^ neutrophil populations expressing different Mmp9 levels (NO, INT, HI) sorted from the WKM of two adult *Tg(lysC:CFPNTR)*^*vi004*^*/Tg(BACmmp9:Citrine-CAAX)*^*vi003*^ zebrafish **(Supplementary Fig. 3A**,**B)**. The generated dataset comprised a total of 18,150 cells passing quality control **(Fig. 3A, Supplementary Fig. 3C-I, Supplementary Table 9)**.

**Fig. 3:**
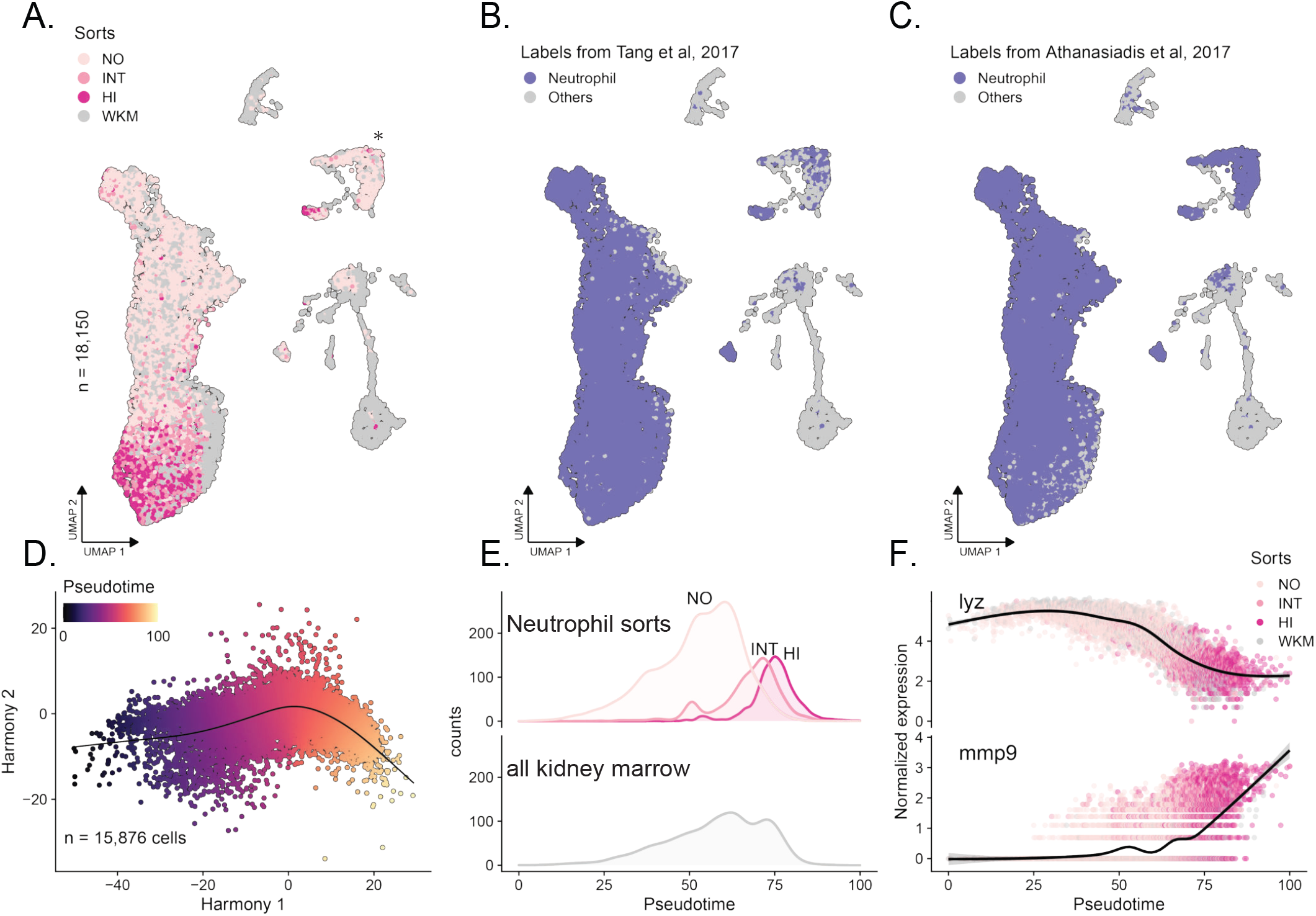
scRNA-seq of zebrafish neutrophils reveals a continuous maturation process. **(A)** Uniform Manifold Approximation and Projection (UMAP) of single-cell RNA-seq data (n = 18,150 cells) showing cells from FACS-sorted neutrophil populations (lysC^+^/mmp9 = NO, INT, HI) and unsorted whole-kidney marrow cells (WKM). **(B-C)** Reference-based labelling of cell types by projecting cells to two zebrafish hematopoietic reference atlases in the same UMAP as in panel A^24, 37^. As expected, a strong overlap with genetically labelled *Tg(mpx:GFP)* neutrophils is observed (highlighted in color). **(D)** Harmony^73^ plot of the inferred neutrophil maturation trajectory in zebrafish (n = 15,876 cells). A continuous trajectory indicating a maturation continuum is observed. **(E)** Line plots of the distribution of sorted (NO, INT, HI) and unsorted neutrophil populations along the inferred trajectory. **(F)** Smoothed expression of *lyz* and *mmp9* in neutrophils along the inferred trajectory.

Sorted neutrophil populations and neutrophils from WKM overlapped in our dataset, indicating that our sorting strategy captured all neutrophils present in WKM **(Fig. 3A)**. To investigate the maturation dynamics, we focused on cells consistently annotated as neutrophils (n = 15,876 cells) based on a bioinformatic mapping to two independent reference datasets^24, 37^ (**Fig. 3B,C**) and used the Slingshot algorithm to infer the structure of the underlying trajectory **(Fig. 3D,E)**^38^. The validity and directionality of this trajectory was supported by (i) the sequential order of the sorted subpopulations along the trajectory (in order: Mmp9^NO^, Mmp9^INT^, Mmp9^HI^; **Fig. 3E)**, (ii) decreasing levels of lysozyme C(*lyz)* and increasing levels of *mmp9* **(Fig. 3F)**, and (iii) a decrease in the number of cycling cells **(Supplementary Fig. 3G)**. A cluster separated from the main neutrophil population was mainly formed by cells in the G2/M phase (**Fig. 3A, Supplementary Fig. 3F**, labelled by *).

We used the tradeSeq^39^ algorithm to define the genes associated with neutrophil maturation (*P*_*adj*_ *< 0*.*05*, top 1500 in descending order based on Wald-statistic; **Supplementary Table 1**). Based on expression patterns of these genes, we partitioned the cells into four maturation phases along the trajectory (P1 = early, P2 = early/intermediate, P3 = intermediate/late, P4 = late). Similarly, we partitioned the genes into three modules (M1 = early, M2 = intermediate, M3 = late) **(Fig. 4A)**. Whereas M1 and M2 foremost cohere to the respective phases, M3 genes are expressed in P3 and P4 **(Fig. 4B)**. Functional enrichment analysis using hypeR^40^ indicated that earlier modules M1 and M2 related to cell cycle and proliferation while module M3 comprised genes associated with functions of mature neutrophils like migration, immune activation, and inflammation **(Fig. 4C, Supplementary Table 3)**.

**Fig. 4:**
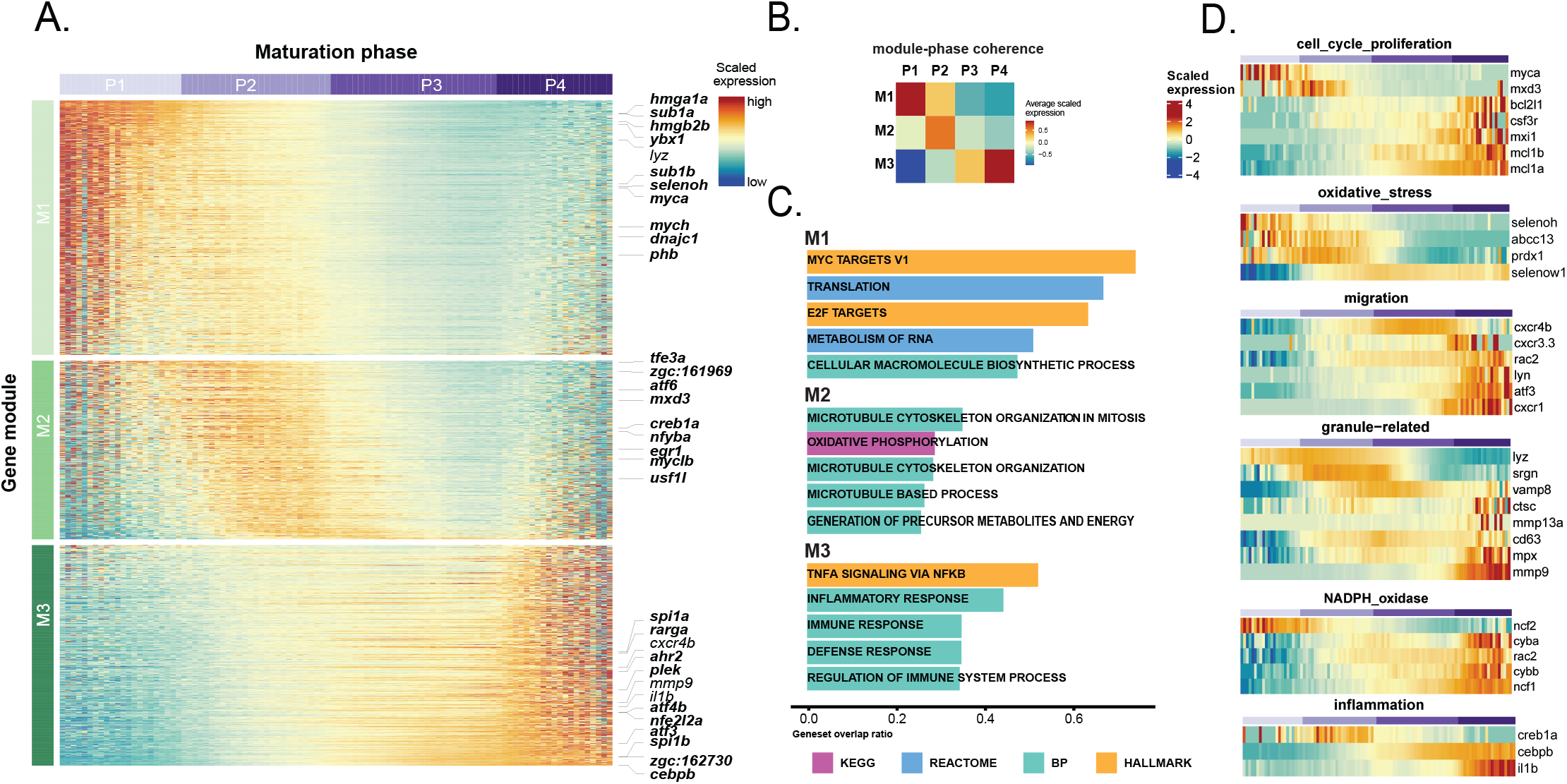
Trajectory analysis uncovers the underlying cell phases and governing gene modules. **(A)** Heatmap showing differentially expressed genes (tradeSeq^39^;P_adj_ < 0.001, top 1500 based on Wald-statistic) along the maturation trajectory. Cells have been allocated to four distinct phases (P1-P4) and into three distinct gene modules (M1-M3) by hierarchical clustering. **(B)** Heatmap summarizing average gene expression per module and phase (from panel A). **(C)** Top-5 enriched gene sets per module from the indicated source databases pathways (hypeR^40^ over-representation analysis). **(D)** Heatmaps of differentially expressed genes (excerpts from panel A) associated with selected neutrophil-related functional pathways selected from the literature.

We next examined genes associated with certain neutrophil functions in detail **(Fig. 4D)**^6, 8, 12, 14^. First, we found proliferation factor *myca* expressed during early maturation (P1), while advancing maturation (P3, P4) was associated with anti-apoptotic genes such as *mcl1a, mcl1b* and *mxi1*, a negative regulator of *myc*^41^. Second, we detected genes related to oxidative stress response (*selenoh*^42^, *abcc13, prdx1*) early in P1, which could aid to protect progenitor cells from oxidative damage. Third, we found a down-regulation of the marrow retention factor *cxcr4b* in P3, and conversely an upregulation of *cxcr1, cxcr3*.*3, lyn*, and the migration-related transcription factor *atf3* in P4, in line with a putative switch towards a migratory phenotype^43^.

Finally, we also found genes encoding granule-related proteins expressed in specific phases **(Fig. 4D)**, these granules have conventionally been used for neutrophil staging^5^. At the beginning of the trajectory *lyz* from P1 on (compare Fig. 3F), primary granule-related genes *cd63, sgrn*, and *mpx* from P2 on, secondary granule-related NAPDH oxidase subunits *cybb* and *cyba* from P3 on, and the gelatinase *mmp9* and *mmp13a*.*1*, a putative MMP1 orthologue, in P4. The expression of these genes is consistent with a staging of human neutrophils from myeloblasts (P1) through pro-myelocytes (P2) and myelocytes/metamyelocytes (P3) to banded/segmented neutrophils (P4). The last phase (P4) was also associated with the upregulation of the pro-inflammatory cytokine *il1b* **(Fig. 4D)**, confirming data from mouse neutrophils^12, 14^.

Taken together, our data demonstrate that neutrophils in the kidney marrow of zebrafish mature in a continuous process advancing from a proliferative stage to a post-mitotic, anti-apoptotic, and migratory phase. Detection of genes associated with certain granule types suggests a similar sequence of granule production as in mammals **(Supplementary Fig. 9)**.

### C/ebpβ governs expression of late granular genes during maturation

Next, we sought to identify the key regulators of the neutrophil maturation process in zebrafish. Many known neutrophil TFs displayed dynamic expression patterns along the trajectory **(Fig. 5A)**^8^. Comparison of transcription factor expression with the expression of genes in each module using dynamic time warping (DTW) analysis highlighted *ybx1*, a splicing factor key in hematopoietic development^44^, *hmgb2b* and *dnajc1* as top regulators of early maturation genes in module M1 (**Fig. 5B-C**; **Supplementary Fig. 4; Supplementary Table 4**,**5**). In contrast, *cebpb* and *atf3* emerged as the top-ranked transcription factors for late maturation (M3). C/EBPβ governs demand-driven granulopoiesis during infection in zebrafish and in mammals^27, 28^ and has also been identified as a late-activated transcription factor in neutrophils in mice ^12, 14^. However, a role during steady-state neutrophil maturation has not been described to date.

**Fig. 5:**
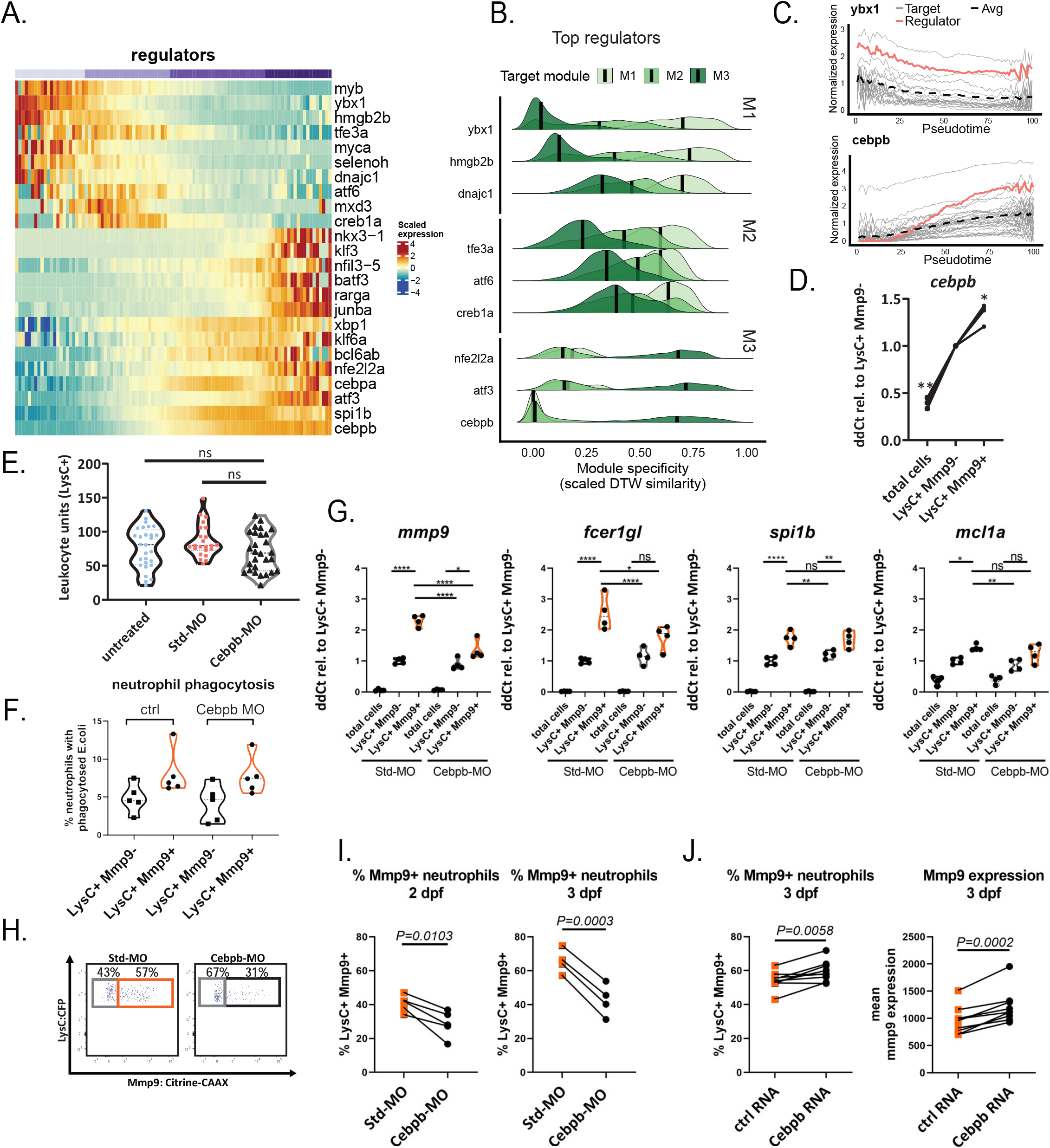
Cebpb is a key regulator of late neutrophil maturation in zebrafish. **(A)** Heatmap showing regulation of selected transcription factors (TFs) during neutrophil maturation. **(B)** Top-3 specific TFs per target module selected by inverse dynamic time warping (DTW) distance (= similarity) between expression of the TF and target genes. **(C)** Expression of top TFs *ybx1*, compared to its putative target genes in module M1 (top), and *cebpb*, compared to genes in M3 (bottom). **(D)** qPCR validation of *cebpb* expression in sorted *lysC:CFP*^*+*^*/mmp9:Citrine*^*+*^ *and lysC:CFP*^*+*^*/mmp9:Citrine*^*-*^ neutrophils. n = 4; repeated measures ANOVA ** p<0.01; * p<0.05; **(E-H & I)** Analyses of Std-morpholino or Cebpb-morpholino treated *Tg(lysC:CFP-NTR)*^vi002^*/Tg(BACmmp9:Citrine-CAAX)*^vi003^ larvae at 3 dpf. **(E)** ImageJ analysis of LysC^+^ leukocyte units from Leica Sp8b confocal images with a HC PL APO CS 10x/0.40 DRY; Zoom 0.85x. **(F)** *in vivo* phagocytosis assay with E.coli-mCherry injected into caudal vein after cebpb Morpholino treatment. **(G)** qPCR of selected target genes of the cebpb regulon on FACS-sorted cells after morpholino treatment. **(H)** Representative flow cytometry blot shows reduction in DP neutrophils. **(I)** Frequencies of LysC^+^Mmp9^+^ neutrophils are reduced after Cebpb morpholino knock-down. Four independent experiments each with pools of ca. 20 larvae per group analyzed by flow cytometry at 2 and 3 dpf after morpholino treatment. Paired t-test. **(J)** Frequencies of LysC^+^Mmp9^+^ neutrophils and mean mmp9 expression are increased after Cebpb overexpression. Analyzed by flow cytometry 3 dpf after mRNA injection (n = 7), paired t-test.

We hypothesized that C/ebpβ is involved in the regulation of neutrophil maturation in zebrafish. After confirming that *cebpb* expression in sorted LysC^+^Mmp9^+^ neutrophils was significantly higher than in LysC^+^Mmp9^-^ neutrophils and unsorted cells (repeated measures ANOVA, *P<0*.*05*, **Fig. 5D**), we targeted *cebpb* translation using a published AUG-binding morpholino. No systemic adverse effects were observed, knockdown did not influence steady-state neutrophil numbers, in line with previous data **(Fig. 5E)**^27^. We examined whether C/ebpβ regulates the augmented phagocytic function of Mmp9^+^ neutrophils, but found no effect on uptake of mCherry-labelled *E*.*coli* **(Fig. 5F)**. *Cebpb* knock down affected expression of the published C/ebpβ target genes^14^ *mmp9* and *fcer1gl* but not of *spi1b* and *mcl1a* in FACS-sorted LysC^+^Mmp9^+^ neutrophils **(Fig. 5G)**^14^. Strikingly, we found that *cebpb* morpholino treatment reduced the frequency of LysC^+^Mmp9^+^ neutrophils at 2 and 3dpf **(Fig. 5H,I)**, indicating an instructive role of C/ebpβ in the differentiation towards mature Mmp9-expressing neutrophils.

Conversely, *cebpb* RNA overexpression led to higher LysC^+^Mmp9^+^ neutrophil frequencies and mmp9 expression levels, further supporting this role **(Fig. 5J)**.

Together, these data indicate that C/ebpβ – in addition to its role in demand-driven hematopoiesis – regulates aspects of steady-state neutrophil maturation such as generation of mature Mmp9^+^ neutrophils, but not their phagocytic function.

### Cross-species comparison identifies conserved gene signatures of neutrophil maturation

Next, we wanted to inquire the grade of conservation of neutrophil maturation trajectories between zebrafish and mammalian species, an important piece of information for investigating and modelling granulopoiesis across species.

Therefore, we compared the inferred neutrophil maturation trajectory from our zebrafish model **(Fig. 6A)** to maturation trajectories based on published mouse and human datasets. We started with a comparison to mouse (scRNA-seq n = 3^10, 12, 14^, **Fig. 6B**, bulk RNA-seq n = 1^11^ **Fig. 6C**) data. For each trajectory, we aligned the expression patterns of genes along the maturation trajectory to the corresponding homolog along the zebrafish trajectory using cross-correlation. We found that gene expression in early development (M1_ortho_) was highly congruent with their mouse orthologues in all datasets. At later stages (M2_ortho_ and M3_ortho_ modules), expression profiles were less consistent but a subset of genes in each module was found conserved across all datasets **(Fig. 6F)**. We expanded the same analysis to human (scRNA-seq n = 3^45, 46, 47^ **Fig. 6D**, bulk RNA-seq n = 1^48^ **Fig. 6E)**, confirming a consistent early expression of M1_ortho_ genes and partial conservation of intermediate and late zebrafish neutrophil maturation signatures in human (M2_ortho_-M3_ortho_). Human neutrophils have been difficult to capture in scRNA-seq assays leading to sparse representation in datasets and under-representation of intermediate maturation phases. We therefore also looked at *in vitro* differentiated neutrophils from human HL-60 promyelocytes, which displayed a very similar expression pattern across all zebrafish gene modules throughout 120 hours of differentiation **(Fig. 6E)**^48^.

**Fig. 6:**
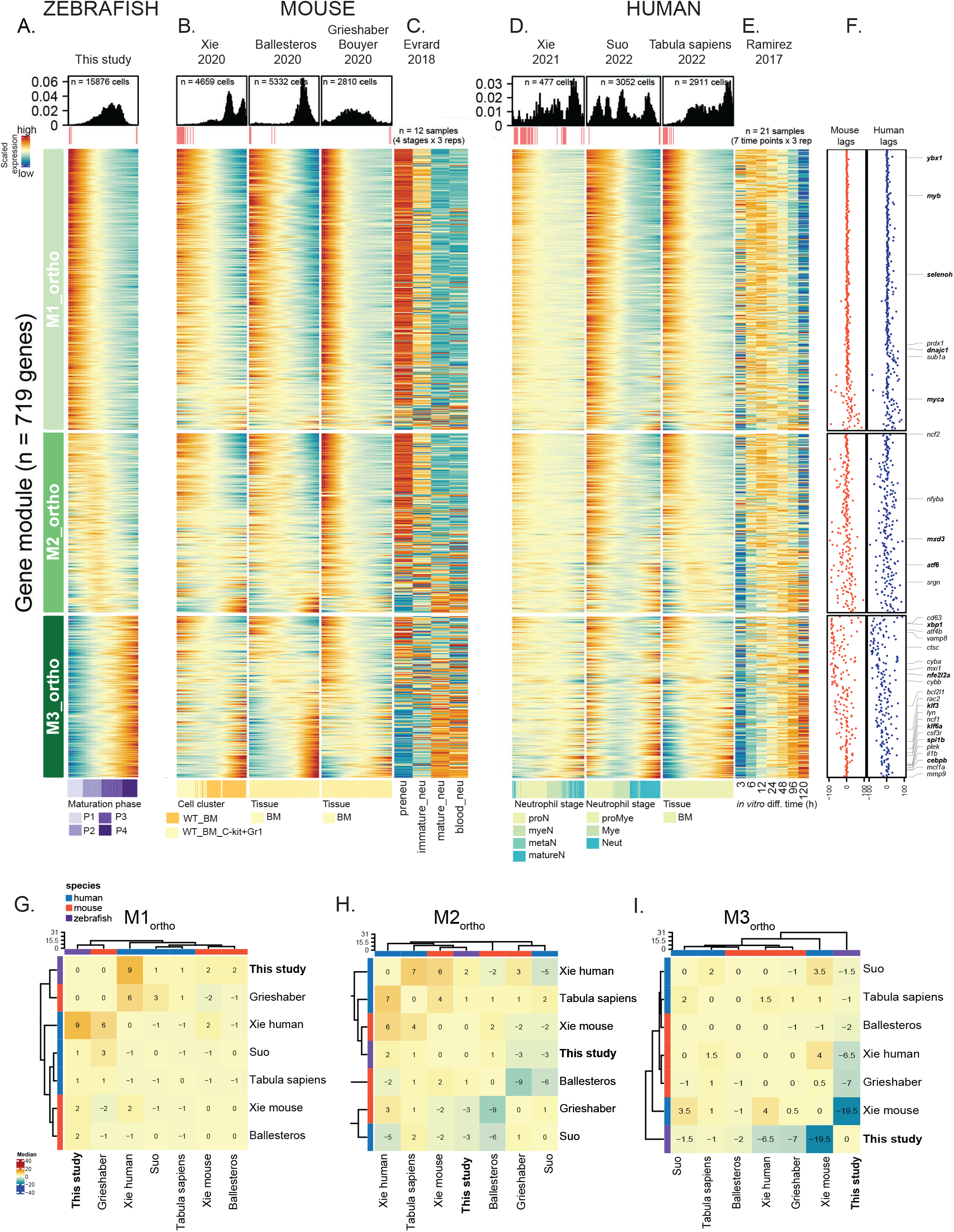
Alignment of expression trajectories across zebrafish, mouse and human reveal concordant and divergent stages of maturation. Cross-species comparison of the orthologues of neutrophil maturation signatures (M1_ortho_, M2_ortho_, M3_ortho_) in zebrafish (heatmap in panel **A**) compared to mouse (**B**,**C**) and human (**D**,**E**) neutrophil datasets^10, 11, 12, 14, 45, 46, 47, 48^. Heatmaps in panels B and D display scRNA-seq data ordered by their own maturation trajectory. Panels C and E show bulk RNA-seq data of FACS-sorted **(C)** and *in vitro* differentiated **(E)** neutrophils in maturation order. In all panels(from top to bottom): top annotations display, dataset annotation, cell density (histograms) and quality-flagged bins (<= 3 cells; potentially less reliable), heatmaps for modules M1_ortho_-M3_ortho_, and author-provided annotations. Left side annotation bars show module membership of each gene orthologue. **(F)** Average cross-correlation lag between mouse and human datasets (from panels **B**,**D**) and the corresponding zebrafish gene. **(G-I)** Heatmaps displaying the median co-phenetic distance of orthologue expression between the indicated datasets, ordered by hierarchical clustering with complete linkage.

We used cross-correlation analysis to quantify the degree of agreement or divergence in gene expression patterns in mouse and human compared to zebrafish. For instance, this analysis confirmed the strong agreement of identified candidate regulators of M1 (*ybx1*) and M3 (*cebpb*) across the three species **(Supplementary Fig. 5A)**. *mmp9* expression lagged behind in zebrafish compared to some human (Tabula Sapiens and Xie 2021) and mouse (Grieshaber− Bouyer 2020) datasets that contain peripheral neutrophils that were not analyzed in our WKM isolates. Conversely, other genes showed species-specific differences; for instance, *tgfbi* was expressed earlier in human but later in mouse trajectories compared to zebrafish. Other genes showed a lag (e.g., *txnipa*) or advance (e.g., *amd1*) compared to zebrafish. To assess the similarity of maturation trajectories at species-level, we calculated the median cross-correlation coefficient per dataset for each module **(Fig. 6F, Supplementary Fig. 5B)** and used hierarchical clustering to group datasets with similar maturation dynamics **(Fig. 6G-I)**. We found that the range of cross-correlation lags for M1_ortho_ and M2_ortho_ was smaller than for M3_ortho_, for which differences between all datasets increased. Taken together, these results indicate a strong conservation (across all species) of early neutrophil maturation, while differences arise at late maturation stages.

Finally, we compared our orthologous gene modules (M1_ortho_-M3_ortho_) to frequently used gene signatures for human immature neutrophils (HAY_BONE_MARROW _IMMATURE_NEUTROPHIL; M39200)^49^. Surprisingly, the majority of putative “immature neutrophil” genes were attributable to late modules (M2_ortho_, M3_ortho_) of zebrafish neutrophil maturation **(Supplementary Fig. 6A)**, but also to late time points of human *in vitro* neutrophil differentiation **(Supplementary Fig. 6B)**, highlighting the necessity for alternative gene signatures of different maturation grades. Our cross-species comparison allowed us to define a pan-species gene signature of neutrophil maturation (absolute cross-correlation lag <= 50; M1_pan_ = 304, M2_pan_ = 176, M3_pan_ = 122; **Supplementary Table 1**,**6**; see *Cross-species integration and comparison* for details).

We asked whether this gene signature could be used to infer the maturation grade of neutrophils in heterogeneous tissues from bulk RNA sequencing data. To this end, we analyzed metastatic neuroblastoma samples^50^. Neuroblastoma is a childhood cancer derived from the sympathetic nervous system that frequently disseminates into the BM. Comparison of BM with (n = 17 datasets) and without tumor cell infiltration (n = 21 controls) thus presents an *in vivo* test case to examine our gene modules. Single-sample Gene Set Enrichment Analysis (ssGSEA)^51^ indicated a differential enrichment of early modules (M1_pan_ and M2_pan_) in control samples opposed to an enrichment of the mature module M3_pan_ in tumor-infiltrated samples **(Fig. 7A, Supplementary Fig. 7A)**. For validation we scored the percentage of segmented neutrophils on BM cytospins from patients with localized (n = 9) and infiltrated (n = 12) neuroblastoma by imaging mass cytometry (IMC; Iridium-intercalator = nuclear; CD15 = granulocytic; **Fig. 7B, C, Supplementary Fig. 7B)**. This confirmed a significant increase of segmented neutrophils in the metastatic group compared to control (Unpaired t-test; *P* = 0.025).

**Fig. 7:**
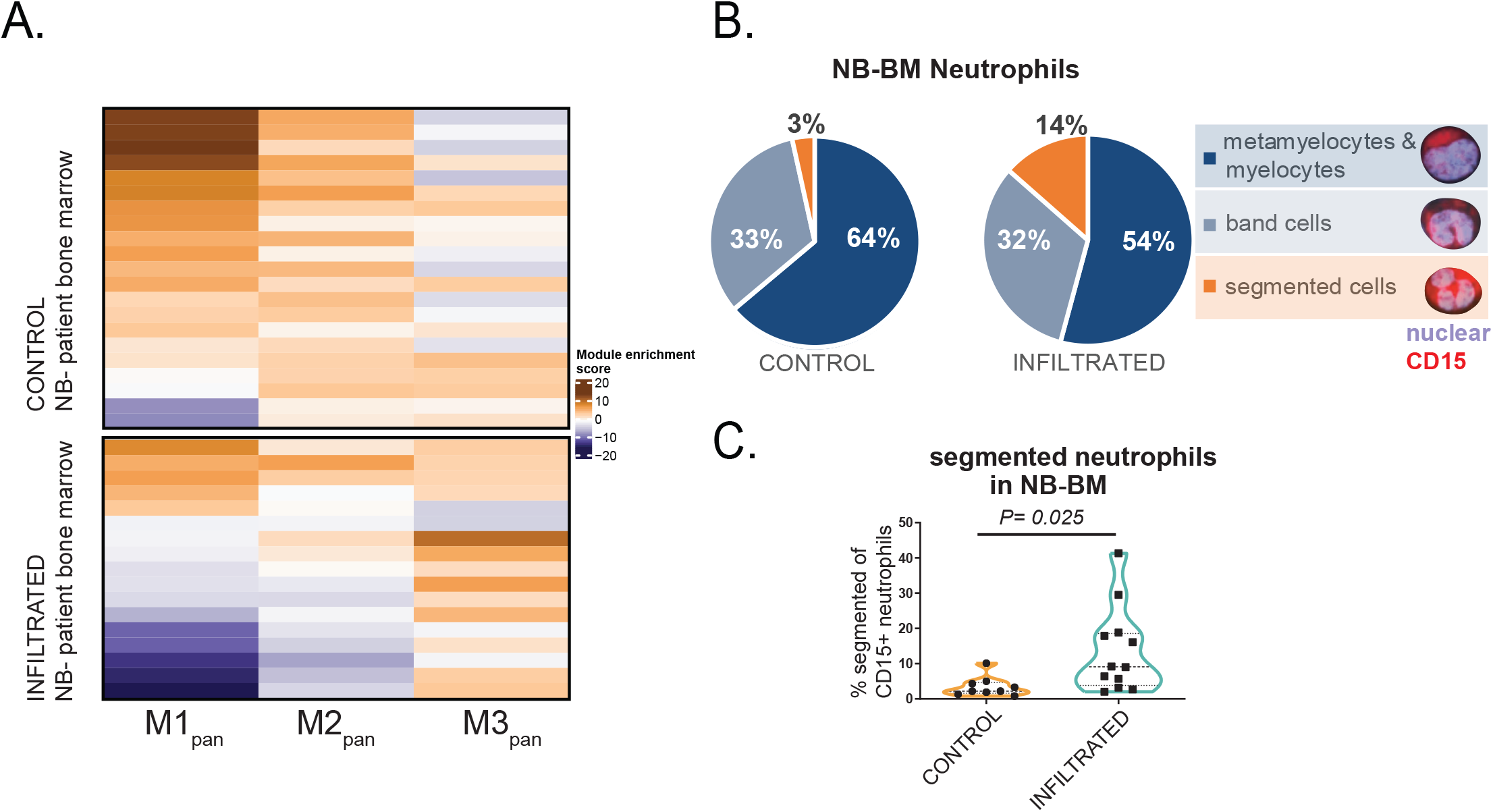
A pan-species neutrophil maturation signature shows the enrichment of mature neutrophils in metastatic BM of neuroblastoma patients. **(A)** Application of the pan-species neutrophil maturation signatures (M1_pan_, M2_pan_, M3_pan_) derived from maturing zebrafish neutrophils (see *Methods* for details on signature definition) on bulk RNA sequencing data from 38 BM samples of patients with metastatic (n = 17, infiltrated) and localized (n = 21, control) neuroblastoma. The heatmap displays the ssGSEA^51^ score of the maturation signatures on RNA-seq samples from 38 neuroblastoma patients with (n =1 7) and without (n = 21) tumor cell bone marrow infiltration. **(B**,**C)** Assessment of neutrophil maturation based on nuclear morphology in BM cytospin samples from patients with metastatic (n = 12, infiltrated) and localized (n = 9, control) neuroblastoma. Samples were stained with DNA-intercalator Iridium and anti-CD15-Bi209, analyzed with IMC and counted blinded. Unpaired t-test.

In summary, we established a comprehensive cross-species transcriptome comparison of neutrophil maturation, suggesting a high degree of conservation between zebrafish and mammalian models. The pan-species gene signature derived from this comparison will present a valuable and robust alternative to existing gene signatures of neutrophil maturation and is applicable for examination of human samples.

## Discussion

In this study, we generated lysC:CFP/mmp9:Citrine transgenic zebrafish, which enabled us to identify and study neutrophils of different maturation grades non-invasively in a living organism. Using this model and single-cell RNA sequencing we generated the first neutrophil maturation atlas in zebrafish as a foundation for a comprehensive cross-species comparison.

In zebrafish, we detected Mmp9 expression in neutrophils (LysC^+^ Mpeg^low^), but rarely in macrophages. This is different from mouse, where in models of both metastatic lung cancer and pancreatic cancer MMP9 was also produced by macrophages^52, 53^. In our model, Mmp9 expression correlated not only with a higher maturation grade but also with classic mature neutrophil effector functions such as phagocytic capability, recruitment towards bacterial infections, and the ability to interact with oncogene-expressing cells. Currently, it is still unclear how these intense interactions between Mmp9^+^ neutrophils and the pre-neoplastic cell clusters are formed. Potentially responsible are integrins upregulated during maturation such as the ones we find in our M3 gene signature, e.g. *itgae*.*2* and *itgb2* (CD18). The latter is well known in leukocyte rolling and important in neutrophil recruitment to sterile wounds^54, 55^. Whether Mmp9^+^ neutrophils and their interactions play a pro- or anti-tumorigenic role needs further investigation.

Our scRNA-seq analysis of neutrophils revealed a continuous neutrophil maturation trajectory similar to observations for mouse neutrophils^12^. While early maturation was controlled by *Cebpe*, the zebrafish orthologue (*cebp1*) did not come up in our signature. Instead, we identified other well-known transcription factors in the early phase such as *myb*^*56*^ and *myca*, as well as y*bx1*, which had previously been implicated in hematopoietic networks^44^. As in mice we found late maturation under the control of *cebpb* and associated with inflammatory pathways and increasing levels of *il1b* and *csf3r*, suggesting a pro-inflammatory state of neutrophils awaiting release from the kidney marrow^12^. In addition to *cebpb*’s known role in emergency granulopoiesis^27, 28^ we found evidence for a role in homeostasis, the expression of tertiary granule genes such as *mmp9* and *fcer1gl*^*57*^, and generation of Mmp9^+^ neutrophils. Our transcriptomic dataset (M1-M3) will be a valuable resource to study neutrophils in zebrafish in homeostasis and under pathological conditions.

To use zebrafish as a model for human disease, it is crucial to delineate similarities and differences between species. The importance of neutrophils as pathogen-fighting cells with their core functions in phagocytosis, ROS production, and degranulation is underlined by their evolutionary conservation. Granular cells are present already in invertebrates such as lancelets (*Branchiostoma*), and myeloperoxidase is even produced by invertebrates^58^. Our comprehensive neutrophil gene expression comparison of zebrafish and mammalian orthologues revealed congruent transcriptional dynamics throughout maturation. The expression of transcription factors *ybx1* and *cebpb* was particularly well synchronized across species, while other genes diverged (e.g., *tgfbi*).

Quantification of similarity allowed us to define a robust pan-species core signature of neutrophil maturation (M1_pan_-M3_pan_), which includes only orthologues with synchronous expression dynamics in zebrafish, mice, and humans. These signatures may help to interpret neutrophil states also in human bulk RNA sequencing data and we validated them here using published data from BM metastases of pediatric neuroblastoma patients^50^. Intriguingly, we detected the mature neutrophil signature (M3_pan_) in patients with disseminated tumor cells. We speculate that this enrichment in mature neutrophils could either result from prolonged retention of mature TANs in the BM or increased recruitment of peripheral neutrophils in response to metastasizing tumor cells. The presence of TANs in BM metastases has been associated with a pro-inflammatory and concurrently immuno-suppressive environment in the BM metastatic niche^50^. In the future, our signature could be applied to relate neutrophil maturation grade to disease progression or outcome.

In conclusion, zebrafish-derived neutrophil maturation modules are conserved and can be translated across species to investigate maturation stages of human neutrophils in metastases samples. The strong homology of neutrophil maturation reinforces the translatability of zebrafish models to study neutrophil biology. The combination of live imaging and transcriptomic approaches in zebrafish will further enable the dissection of the role of immature and mature neutrophils in the tumor microenvironment and the consequences of their interactions with tumor cells.

## Acknowledgements

We would like to thank Dieter Printz for FACS sorting and the Biomedical Sequencing Facility of the CeMM Research Center for Molecular Medicine of the Austrian Academy of Sciences for next-generation sequencing. We would like to acknowledge Cristina Santoriello and Marina Mione for the kind gift of the *HRAS_G12V:UAS:CFP* plasmid. We would like to acknowledge the CCRI Biobank and Marie Bernkopf. Peter Repiscak for providing support on pre-processing of NB-BM sequencing data. We would like to thank the members of the Innovative Cancer Models group and Dietmar Herndler-Brandstetter for their insightful comments on the manuscript. This study was supported by the St. Anna Kinderkrebsforschung (to S.T.M., R.L., F.H., and M.D.), the Austrian Science Fund (FWF) through grants I4162 (ERA-NET/Transcan-2 LIQUIDHOPE; to S.T.) and P35841-B (MAPMET; to S.T.) and the Swiss Government Excellence Scholarship (to D.L.) and the EC H2020 grant no. 826494 (PRIMAGE; to R.L.).

## Authorship contributions

SK designed and performed experiments, analyzed data and wrote the manuscript. MS performed bioinformatics analysis including collection and preprocessing of public data, and wrote the manuscript. DL performed IMC on neuroblastoma BM samples. KF supported BAC screening. LS performed the 10x Genomics workflow. FR designed the graphical abstract. FR and EB developed new imaging methods. RL has provided patient samples and clinical data. BB provided access to IMC instruments. FN, TL and DT provided reagents. MF supervised scRNA-seq analysis. STM and BB conceptualized and supervised the analysis of human BM samples. FH and MD designed and analyzed experiments, supervised all work, and wrote the manuscript. All authors have contributed to writing the manuscript.

## Methods

### Zebrafish care and zebrafish transgenic lines

Zebrafish experiments were performed under research licenses GZ:565304/2014/6 and GZ:534619/2014/4 according to the guidelines of the local Austrian authorities (Vienna Magistrat MA58). Zebrafish (*Danio rerio*) were kept under standard conditions^59^ in a research fish facility (Tecniplast, Italy). Larvae were kept in egg medium with 20 mg/l phenylthiourea (Merck) from 22 hpf to avoid pigmentation. Tricaine was used as an anesthetic.

The following transgenic lines were used *Tg(lysC:CFP-NTR)*^*vi002*^, *Tg(lysC:dsRed)*^*nz50Tg*^, *Tg(mpeg1:mCherry)gl23, Et(kita:GAL4)*^*hzm1*^, *Tg(UAS:EGFP-HRAS_G12V)*^*io006*^, *Tg(HRAS_G12V:UAS:CFP)*^*vi004*^, *Tg(BACmmp9:Citrine-CAAX)*^*vi003*^. *Tg(BACmmp9:Citrine-CAAX)*^*vi003*^ fish (abbreviated mmp9:Citrine) were generated by BAC transgenesis according to published protocols^60, 61^. In short: Identity of annotated BAC CH211-269M15 (105.8 kb) containing mmp9 gene and regulatory regions (BACPAC Resources) was confirmed by sequencing, recombineered with iTol2 sites and with membrane-targeted fluorescent Citrine-CAAX DNA inserted at the start codon of *mmp9*. BAC DNA (67 ng/µl) was micro-injected into fertilized zebrafish eggs together with Tol2 transposase mRNA as previously described^62^. pDESTlysC:CFPNTR (#54) was generated by gateway recombination of p5lys:C (#31), pENTR1A-CFPNTR (#51) and pDESTTol2pA2 (kind gift of Chi-Bin Chien). *Tg(HRAS_G12V:UAS:CFP)* plasmid was constructed and kindly provided by the Mione lab. The transgenesis constructs were injected into fertilized zebrafish eggs at 25 ng/µl together with 25 ng/µl Tol2 RNA. Injected zebrafish were grown up to adulthood and screened for germline transmission.

### Flow cytometry and FACS

Single-cell suspensions from adult zebrafish (3 months) spleens or kidneys were prepared by mashing or pipetting, respectively, and pipetting through a cell strainer. To isolate cells from larvae, they were immersed in 10 mM DTT (Merck) in E3 to remove mucus, then digested with Liberase Blendzyme TM at 1.1 U/ml and Dnase I at 40 μg/ml (Merck) in HBSS under constant shaking at 37°C for 40 minutes. Samples were run on an LSRFortessa cytometer (Becton-Dickinson) and analyzed in FlowJo or sorted using a FACSAria Fusion.

### Injection of bacteria and phagocytosis assay

*E. coli* labelled with mCherry (pZS*12-mCherry-KANr: PRlambda-mCherry in pZSstar, SC101* ORI)^63^ were injected at OD = 2 into the caudal vene area or otic vesicle as previously described^64^. To study the phagocytic capacity of neutrophils, cells from *Tg(lysC:CFP-NTR)*^*vi002*^*/ Tg(BACmmp9:Citrine-CAAX)*^*vi003*^larvae were analyzed after 6 hpi by flow cytometry as described above or imaged using confocal microscopy.

### Morpholino and mRNA micro-injection

We used a previously published C/ebp-β translation blocking morpholino (5′- GATCTTAACACCCGCCGGATTGCG-3′) and a negative control morpholino (5′- CCTCTTACCTCAGTTACAATTTATA-3′) from Gene Tools LCC^27, 65^. Efficacious doses (1 pmole and 0.5 pmole, respectively) were determined empirically. Full-length *cebpb* mRNA (50 pg), prepared by mMessage mMachine T7 Ultra transcription (ThermoFisher) or morpholinos were injected into one to two-cell stage embyos from *Tg(lysC:CFP-NTR)*^*vi002*^*x Tg(BACmmp9:Citrine-CAAX)*^*vi003*^ crosses.

### Cytospin

Kidney marrow cells of adult *Tg(lysC:dsRed)*^*nz50Tg*^*/ Tg(BACmmp9:Citrine-CAAX)*^*vi003*^ fish (aged three months) were FACS sorted into Mmp9^+^ LysC^+^ or Mmp9^-^ LysC^+^ fractions and spun in a cytocentrifuge (Fisher Scientific) onto slides according to the manufacturer’s instructions and stained using Pappenheim solution (Merck).

### IMC analysis of neuroblastoma BM cytospin preparations

Archived cytospin preparations of human BM aspirates from patients with localized and metastatic neuroblastoma have been obtained after institutional review board approval and informed consent by patients or their guardians from the CCRI Biobank (EK1853/2016, EK1224/2020). Samples were thawed for 15 min at RT, washed twice in TBS, fixed in 4% PFA (Carl Roth) at 4°C for 30 min, washed again twice in TBS and then blocked with 3% BSA (Carl Roth) and 0.1% Tween-20 (Merck) in TBS at RT for 1 h. Samples were incubated overnight at 4°C with anti-CD15(HI90)-Bi209 (primary antibody purchased from Biolegend; conjugated with Fluidigm’s MaxPar labeling kit) diluted at 5 µg/ml in 0.1% Tween-20 in TBS. On the following day, samples were stained with the DNA intercalator Iridium (Fluidigm) and then dried with pressured air before IMC measurement with the Hyperion imaging system (Fluidigm). Morphological assessment was subsequently performed in the image analysis software QuPath^66^.

Nuclear morphology of CD15^+^ cells was assessed and cells were classified as segmented cells, band cells or metamyelocytes and myelocytes (3 ROI/patient; n = 248-330 cells/patient). Cells were counted using the QuPath counting tool.

### Quantitative real-time PCR

RNA from FACS sorted cells was prepared using RNeasy Micro kit (Qiagen) and transcribed with High Capacity cDNA transcription kit. qPCR was performed using Maxima SYBR green mix on a 7500 Fast real-time PCR machine (all Applied Biosystems). **Primers used:**

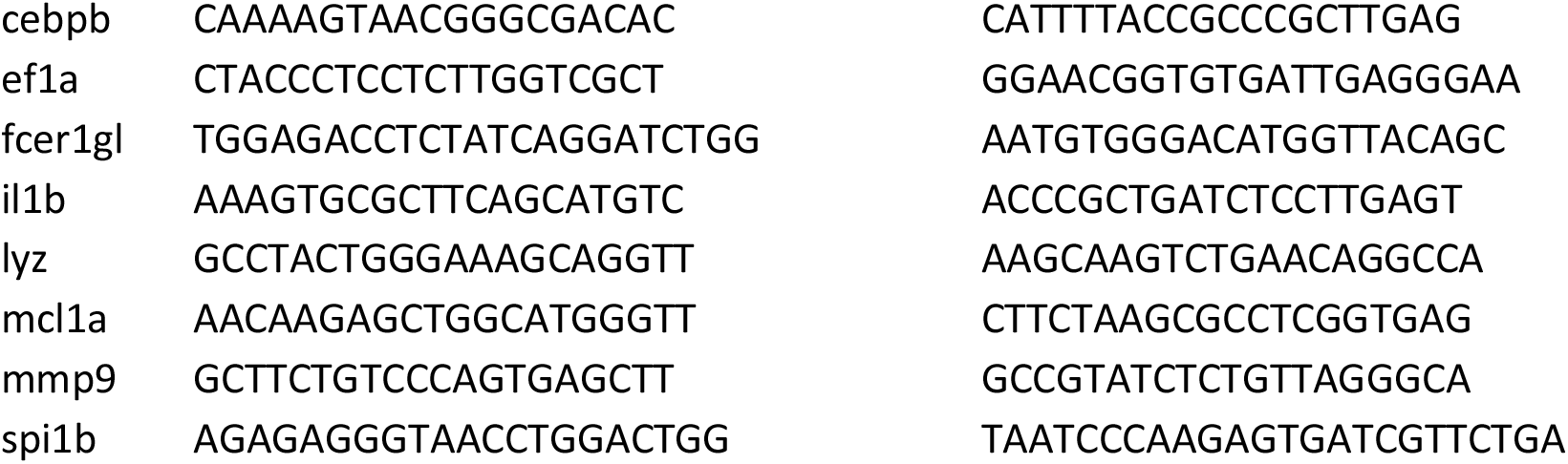

### Imaging

For imaging larvae were pre-treated in E3/PTU, anaesthetized with 0.02% tricaine and embedded in 1.2% low-melting agarose (Merck) on glass bottom dishes (D35-14-1.5-NJ, Cellvis, USA) as described previously^67^. Images were rendered using Leica LAS software, Photoshop CS6 (Adobe) and Fiji ImageJ, e.g. for preparing cell tracks. Leukocyte units were analyzed according to^68^.

### Single-cell RNA sequencing

Kidney marrow from 2 adult, six-month old, male fish was isolated, labeled using lipid-tagged indices as published (MULTI-seq)^69^. In short: each kidney marrow was split into four portions, labelled with lipid anchor plus individual barcode solution (2 μM) for 5’ on ice and then incubated with lipid co-anchor (2 μM) for 5’ on ice. Each cell portion was individually FACS-sorted to obtain one population (mmp9 NO, INT, HI or WKM). All cells were gated on live gate, for WKM debris was excluded in a FSC/SSC gate, and mmp9 NO, INT, HI were gated on LysC:CFP positivity and different levels of Mmp9:Citrine-CAAX expression **(Suppl. Fig 3B)**. For multiplexing, the two WKM populations were sorted into one well (A2) and processed together (20,000 cells each; total: 40,000). Similarly, the six LysC^+^ populations were sorted into one well (A1) and processed together (total: 41,000 cells). Here, cell numbers were ranging from 1,700 to 10,000 cells/population.

Single cell suspensions were immediately subjected to scRNA-seq using the Chromium Single Cell Controller and Single Cell 3’ Library & Gel Bead Kit v3.1 (10x Genomics, Pleasanton, CA), according to the manufacturer’s protocols (10x Genomics) and sequenced by protocol. Sequencing was performed at the Biomedical Sequencing Facility of the CeMM Research Center for Molecular Medicine of the Austrian Academy of Sciences (Vienna, Austria) using the Illumina NovaSeq platform and the 50bp paired-end configuration with adapted read-lengths for both forward and reverse reads.

### Bioinformatics analyses

#### Read processing, quality control, and normalization

We used the CellRanger v3.1.0 software (10x Genomics) for cell demultiplexing and alignment to GRCz11-3.1.0 zebrafish reference transcriptome that had been expanded to include the sequences of reporter genes (Citrine and CFPNTR, sequences from snapgene.com). The R statistics software v4.0.3 was used to carry out the entire analysis workflow. Processed data was loaded into R for sample demultiplexing (package deMULTIplex^69^ v1.0.2; cell classification step was stopped when negative cell number dropped below 100). Only cells classified as singlets or negatives were retained, while doublets were excluded. Next, we loaded the counts into Seurat v4.0.2^70^ and performed quality control by only including features detected in at least 20 cells (n = 12,619 out of 25,109 features retained), and cells with a minimum of 500 features, mitochondrial reads proportion less than 10%, and doublet score, calculated using function *doubletCells* (package scran^71^ v1.18.3;default parameters), below 3 (n = 19,373 out of 28,534cells28534 cells retained). Remaining negative cells (that is, cells without an assigned sample label; n = 4,723) from each sequencing run were reclassified separately using Linear Discriminate Analysis (function *lda* from the package MASS v7.3-53; default parameters), only relabeling negative cells with a posterior probability >= 0.95 (n = 3,500 cells), yielding a final dataset of 18,150 cells. Next, we normalized raw read counts using SCTransform^72^ v0.3.2 and integrated using harmony v1.0^73^ (default parameters) the batch effect of different sequencing runs and fish. We performed low-dimensional projection using UMAP based on the top 30 Harmony components. Reference-based cell annotation using Seurat^70^ v4.0.2 was carried out by mapping our data to two atlases of hematopoiesis in zebrafish^24, 37^ (preprocessing as described above) according to the workflow recommended by the developers (https://satijalab.org/seurat/articles/multimodal_reference_mapping.html; accessed on 18-Nov-2022). Cells identified as neutrophils in both atlases (n = 15876) were selected for downstream analysis. Finally, after subsetting neutrophils, we excluded hemoglobin-related genes (based on the following regular expression vector “^hb[ba]|si:ch211.5k11.8”) and features detected in less than 20 neutrophils to minimize the effect of other cell types on the downstream analysis of neutrophils.

#### Inference of neutrophil maturation trajectories, associated genes, and top regulators

Continuous maturation trajectories were inferred using slingshot^38^ v1.8.0 (default parameters) based on the top two Harmony components of the dataset^73^. We then discretized the trajectory into 100 bins by averaging overlapping cells at each bin. We used the function *fitGAM* (package tradeSeq^39^ v1.4.0, parameters: knots = 6, cellWeights = rep(1, #genes)) to carry out differential expression analysis along the inferred trajectory using pseudotime values from slingshot. We excluded mitochondrial and ribosomal genes, as well as genes with less than 5 cells with 3 or more reads. Genes that passed the cutoff (P_adj_ < 0.05) were put in descending order based on Wald-statistic and the top 1500 genes were selected as maturation-associated genes for further analysis (**Supplementary Table 1**). To define gene modules and maturation phases, we used the binned expression of these maturation-associated genes, calculated the dynamic time warping (DTW) distance using function *dist* (package proxy v0.4-25; method = “dtw”), and used hierarchical clustering (parameters: method = ward.D2”). The resulting dendrograms were cut into three gene modules and four maturation phases, which was the optimal number based on the lowest Kelley-Gardner-Sutcliffe penalty using function *kgs* (package maptree v1.4-7; default parameters) (**Supplementary Fig. 8**). To interpret and characterize the genes in each maturation module, we used the function *hyper* (package hyper^*40*^ v2.0.1; default parameters) with five datasets (CP:KEGG, GO:BP, CP:WIKIPATHWAYS, CP:REACTOME, HALLMARK) retrieved from Molecular Signatures Database (MSigDB) using *msigdbr* function (package msigdbr v7.5.1; species = “Danio rerio”), to carry out over-representation analysis (fdr < 0.1; **Supplementary Table 3**), selected the top five genesets with lowest FDR, and plotted their ratio of overlap with the gene modules. To identify the transcription factors (“top regulators”) that best explain the variability in the expression patterns of each gene module, we collected a list of zebrafish transcription factors based on both AnimalTFDB3 database (date of retrieval: 21 June 2021)^74^ and manual selection from literature^6, 8, 12, 14^, and calculated the DTW distance between each transcription factor and all target genes^74^. DTW allows to account for the expected lag between transcription factor expression and target gene activation. Next, we transformed the DTW distance *d*_*il*_ for gene *i* and transcription factor *l* into a “scaled DTW similarity” *s*_*il*_, as follows: *s*_*il*_ *= 1 – (d*_*il*_ */ max(d*_*i*_)). Finally, to rank putative regulators per module, we used the function *dunn_test* (package rstatix v0.6.0; default parameters) to run post-hoc test following Kruskal-Wallis test of transcription factor-target gene pairs similarities across the three modules (**Supplementary Table 5**).

#### Cross-species integration and comparison

For cross-species comparisons, we obtained RNA-seq and scRNA-seq read counts from multiple published studies^10, 11, 12, 14, 45, 46, 47, 48^ (see **Data Availability** section).

Bulk RNA-seq data were normalized for library size and multiplied by a scaling factor of 1e6 before log_2_-transformation. For scRNA-seq data, we preprocessed the data following the same approach described under *Read processing, quality control, and normalization* section, with the following exceptions: We did not carry out further quality control filtration on data from Suo 2022^47^ (minimum of 501 features and 2001 UMI counts per cell) and Tabula Sapiens^45^ (minimum of 200 features and 2500 UMI counts per cell).

In the case of neutrotime^12^ model, we used the pre-defined embedding provided by the authors via ImmGen single cell explorer (https://singlecell.broadinstitute.org/single_cell/study/SCP1019/ly6-neutrophils-from-bone-marrow-blood-spleen?scpbr=immunological-genome-project#study-download)

We then selected only healthy bone-marrow neutrophils (using provided cell annotations) (**Supplementary Table 8**). Subsequently, we carried out trajectory inference using slingshot based on the top two PCA components and discretization into 100 bins, as explained above. Next, we performed homology mapping across zebrafish, mouse, and human using *getLDS* function (package biomaRt v2.46.3; default parameters). Common genes from human and mouse were renamed to the corresponding homologous zebrafish gene using a modified version of *RenameGenesSeurat* function (package Seurat.utils v1.4.7; https://doi.org/10.5281/zenodo.7228243). Cases of 1-to-many homology were resolved by mapping the genes to the most expressed homolog based on the attribute “detection_rate” from the sctransform model. We kept only the set of genes that are common across all datasets, scaled each of the datasets separately, and ordered the genes based on their expression pattern across all datasets using the *seriate* function (seriation 1.3.1; method = “PCA”) and the corresponding module. Finally, we used cross-correlation to align the gene expression pattern along maturation trajectory in mouse and human datasets onto zebrafish using the *ccf* function (stats v4.0.3; default parameters) and recorded the mean lag at which highest cross-correlation was achieved among mouse and human datasets per gene (**Supplementary Table 7**).

The pan-species neutrophil maturation signatures (M1_pan_, M2_pan_, M3_pan_) were defined based on two inclusion criteria: First, consistent expression pattern between human and zebrafish (|maximum cross-correlation lag| <= 50). Second, we excluded unique known markers of other cell types (Additional file 5 from xCell^75^). We then obtained normalized count data from neuroblastoma bulk RNA-seq^50^ and used this signature to carry out single-sample gene set enrichment analysis (ssGSEA) using function *ssgsea* (package corto^51^ v1.1.11 ;default parameters).

## Data availability

Single-cell RNA sequencing data will be deposited in the Gene Expression Omnibus (GEO) prior to publication. We made use of the following publicly available datasets in our study:

- Single-cell RNA-seq: E-GEOD-100911, E-MTAB-5530, GSE137539, GSE165276, GSE149938, GSE142754, [Myeloid cells from the fetal immune atlas](https://developmental.cellatlas.io/fetal-immune), [Tabula Sapiens -Immune](https://cellxgene.cziscience.com/collections/e5f58829-1a66-40b5-a624-9046778e74f5)
- Bulk RNA-seq: GSE79044, GSE109467, neuroblastoma data was provided by the authors. Others: [neutrotime model](https://singlecell.broadinstitute.org/single_cell/study/SCP1019/ly6-neutrophils-from-bone-marrow-blood-spleen?scpbr=immunological-genome-project), [AnimalTFDB3.0-zebrfish](http://bioinfo.life.hust.edu.cn/static/AnimalTFDB3/download/Danio_rerio_TF), [xCell signatures Additional file 5](https://static-content.springer.com/esm/art%3A10.1186%2Fs13059-017-1349-1/MediaObjects/13059_2017_1349_MOESM5_ESM.xlsx), additional metadata of GSE109467 was kindly provided by the authors.

## Code availability

Computer code used for the data analysis in this paper will be shared via our GitHub page (https://github.com/cancerbits).

