## Supplementary figures for "Cross-species analysis identifies conserved transcriptional mechanisms of neutrophil maturation"

**Extended Data Fig. 1: Newly generated BAC transgenic line *Tg(BACmmp9:Citrine-CAAX)$^{vi003}$* reporting epithelial and leukocyte expression. (A)** *Tg(BACmmp9:Citrine-CAAX)$^{vi003}$* zebrafish exhibit transcriptional activity of the mmp9 locus in epithelial cells of the skin and gut and in dispersed cells. F1 mmp9:Citrine larvae were analyzed using fluorescence confocal microscopy on a Leica Sp8 with a HCX PL APO CS 10x/0.40 DRY objective for Citrine (in yellow) at 2 and 5 dpf. Scale bars 250 µm. Insets show expression in the epithelia of the distal gut and the tail fin. Brightfield (bottom left) and fluorescence pictures of adult fish (3 months) taken with a Zeiss Axio Zoom.V16 stereo with a PlanNeoFluarZ 1x/0.25 FWD 56 mm using an Axiocam 503 color camera show Citrine (shown in green, due to filter) expression in the region of the anus. **(B)** Mmp9 RNA is enriched in whole kidney marrow cells from *Tg(lysC:CFP-NTR)$^{vi002}$/Tg(BACmmp9:Citrine-CAAX)$^{vi003}$* double transgenic adult fish sorted for Citrine expression. Kidneys were isolated and FACS sorted into CFP+ Citrine- or CFP+ Citrine+ cells, which were analyzed by quantitative real-time PCR with primers for mmp9. N = 3, paired t-test *P* = 0.0144.

**Extended Data Fig. 2: Immune activation regulates frequency of mmp9+ neutrophils**

**(A)** Representative flow cytometry histograms showing induction of *mmp9:Citrine* expression in neutrophils gated on *lysC:CFP$^+$* by wounding through PBS injection (left) or administration of bacteria (right). **(B)** Graph summarizing results of six independent activation experiments analyzed by flow cytometry indicating an increased frequency of Mmp9$^+$ cells within the *lysC:CFP$^+$* neutrophil population upon *E. coli* injection. Paired t-test; *P* = 0.009 **(C)** Upregulation of *mmp9:Citrine* in neutrophils gated on *lysC:CFP$^+$* in the kita pre-neoplastic skin lesion model *Et(kita:GAL4)$^{hzm1}$x Tg(HRAS_G12V:UAS:CFP)$^{vi004}$* at 3 dpf. Graph summarizing results of three independent tumor experiments analyzed by flow cytometry. Paired t-test; *P* = 0.005

**Extended Data Fig. 3: Workflow and quality control of scRNA-seq of zebrafish kidney marrow neutrophils (A)** Schemata showing workflow for cell isolation, multiplexing and scRNA-seq. **(B)** FACS gating strategy for neutrophil isolation for 10x Genomics analysis. WKM cells were sorted on a FACS Aria II gating on live (7-AAD negative), FSC/SSC gate to exclude debris and dead cells. Neutrophils were gated on *lys:CFP+* and different levels of *Mmp9:Citrine-CAAX* expression. UMAP representation of cells labelled by biological (**C**, Fish) and technical covariates (**D**, Pool = FACS well; **E,** Batch), inferred cell cycle phase **(F)**, proportion of mitochondrial reads **(H)**, and $log_{10}$ raw UMI counts **(I)**. **(G)** Bar plot showing the proportion of each cell cycle phase per sorted subset. **(J)** Stacked bar plot showing the number of neutrophils in the sorted and unsorted WKM subsets, along with the relative frequency of cells from neutrophil-major and neutrophil minor clusters displayed on a small UMAP in the same plot.

**Extended Data Fig. 4: Identification of candidate regulators per gene module by dynamic time warping analysis**

Ridge plots showing the distribution of transcription factor specificity for each gene across the three modules. Y-axis show all transcription factors grouped by the corresponding module. The X-axis indicates the specificity score of a transcription factor with respect to the genes of each module. Briefly, specificity measures the similarity of a target gene expression pattern to a candidate regulator, compared to all other transcription factors (see *Methods* for the formula).

**Extended Data Fig. 5: Cross-correlation analysis of gene expression during zebrafish and mammalian neutrophil maturation**

**(A)** Line plots showing seven selected examples of genes with different cross correlation patterns. *ybx1* shows a good concordance across species with highest correlation centered at zero lags from zebrafish. *mmp9* expression shows a noticeable lag to an earlier maturation phase relative to zebrafish with cross correlation peaking at negative lags. *tgfbi* shows contradictory pattern between mouse and human. **(B)** Cross-correlation lags for all commonly differentially expressed genes in all datasets. A positive value indicates a gene is expressed at a later stage relative to its expression stage in zebrafish, and *vice versa*.

**Extended Data Fig. 6**: **HAY_immature signature pre-dominantly includes late neutrophil maturation genes**. **(A)** Evaluation of the overlap between zebrafish gene modules and genes from HAY_immature signature[49]. **(B)** Heatmap analysis of HAY_immature signature on Ramirez *et a*l.[48] *in vitro* differentiated neutrophils.

**Extended Data Fig. 7: Human Neuroblastoma-BM RNA-seq analysis. (A)** Scaled ssGSEA[51] enrichment scores for $M1_{pan}$, $M2_{pan}$, and $M3_{pan}$ genes in bulk RNA-seq data[50] from 38 BM of patients with metastatic (n = 17, infiltrated) and localized (n = 21, control) neuroblastoma. Line plots show scores for each dataset. **(B)** Frequency of myelocytes and metamyelocytes, band cells and segmented neutrophils among all $CD15^+$ neutrophils in BM cytospin samples from patients with metastatic (n = 12, infiltrated) and localized (n = 9, control) neuroblastoma. Samples were stained with Iridium and anti-CD15-Bi209, analyzed by IMC and assessed for their morphology in QuPath. Line plots show frequencies of each neutrophil subtype per patient.

**Extended Data Fig. 8: Optimal selection of the number of phases and modules.** Using Kelley-Gardner-Sutcliffe penalty for optimal pruning of the hierarchical cluster tree**.** Minimum penalty corresponds to the suggested number of clusters.

**Extended Data Fig. 9: Model for neutrophil development in zebrafish based on scRNA sequencing.**

Association of gene expression with modules and phases as determined by scRNA-seq. Granule stage and morphology were assumed by comparing gene expression (as in Figure 4D, Supplementary table 2) with what was previously published for neutrophils.

**Extended Data Fig. 10: Graphical abstract**

**Movie 1: Recruitment of Mmp9$^+$ cells to wound.** Behavior and recruitment of neutrophil subpopulations with different mmp9:Citrine levels (*lysC:CFP$^+$/mmp9:Citrine$^+$ and lysC:CFP$^+$/mmp9:Citrine$^-$* ) and macrophages (*mpeg:mcherry$^+$*) to a needle inflicted wound in triple transgenic *Tg(lysC:CFP-NTR)$^{vi002}$/Tg(BACmmp9:Citrine-CAAX)$^{vi003}$/Tg(mpeg1:mCherry)$^{gl23}$* zebrafish larvae at 3 dpf imaged from around 20 min post- injury. Time-lapse maximum projections. Z stacks were acquired every 50 s at 3 µm intervals, with a HC PL APO CS2 40x/1.10 WATER objective on a Leica Sp8 confocal microscope using LAS X software, Zoom: 1.2 x. Scale bar = 25µm

**Movie 2: Migration behavior of an Mmp9$^+$ neutrophil showing projections around a cluster of transformed kita/RAS cells**. Representative time-lapse maximum projections acquired in quadruple transgenic *Et(kita:GAL4)$^{hzm1}$/Tg(UAS:EGFP-HRAS_G12V)$^{io006}$/Tg(lysC:CFP-NTR)$^{vi002}$/Tg(BACmmp9:Citrine-CAAX)$^{vi003}$* zebrafish larvae starting at 102 hpf with a 40x objective. Z stacks were acquired every 51 s at 2 µm intervals with a HC PL APO CS2 40x/1.10 WATER objective on a Leica Sp8 confocal microscope using LAS X software, Zoom: 2 x. Scale bar = 25 µm

**Movie 3: Interactions of Mmp9$^+$ and Mmp9$^-$ neutrophils with transformed kita/RAS cells.**
Representative time-lapse maximum projections acquired in quadruple transgenic *Et(kita:GAL4)$^{hzm1}$/Tg(UAS:EGFP-HRAS_G12V)$^{io006}$/Tg(lysC:CFP-NTR)$^{vi002}$/Tg(BACmmp9:Citrine-CAAX)$^{vi003}$* zebrafish larvae starting at 78 hpf. Z stacks were acquired every 3 min 39s at 3 µm

intervals with a HC PL APO CS2 40x/1.10 WATER objective on a Leica Sp8 confocal microscope using LAS X software, Zoom: 1.7 x. Scale bar = 50µm

**Tables:**

**Supplementary table 1: Maturation-associated genes differential expression statistics and module assignment.**

**Supplementary table 2: Gene expression of maturation-associated genes along the discretized maturation trajectory**

**Supplementary table 3: Output of overrepresentation analysis of gene-modules based on msigdb databases.**

**Supplementary table 4: DTW distance and similarity for all transcription factor – target gene pairs.**

**Supplementary table 5: Output of Dunn's test of transcription factors similarities to target genes across modules.**

**Supplementary table 6: Human and mouse orthologous utilized in mapping gene across datasets in cross-species heatmap.**

**Supplementary table 7: Output of cross-correlation analysis for each dataset separately and aggregated within species.**

**Supplementary table 8: Overview of utilized external datasets.**

**Supplementary table 9: Summary of CellRanger metrics for the sequencing runs.**

Supplementary figure 1:
Newly generated BAC transgenic line Tg(BACmmp9:Citrine-CAAX)[vi003] reporting epithelial and leukocyte expression.

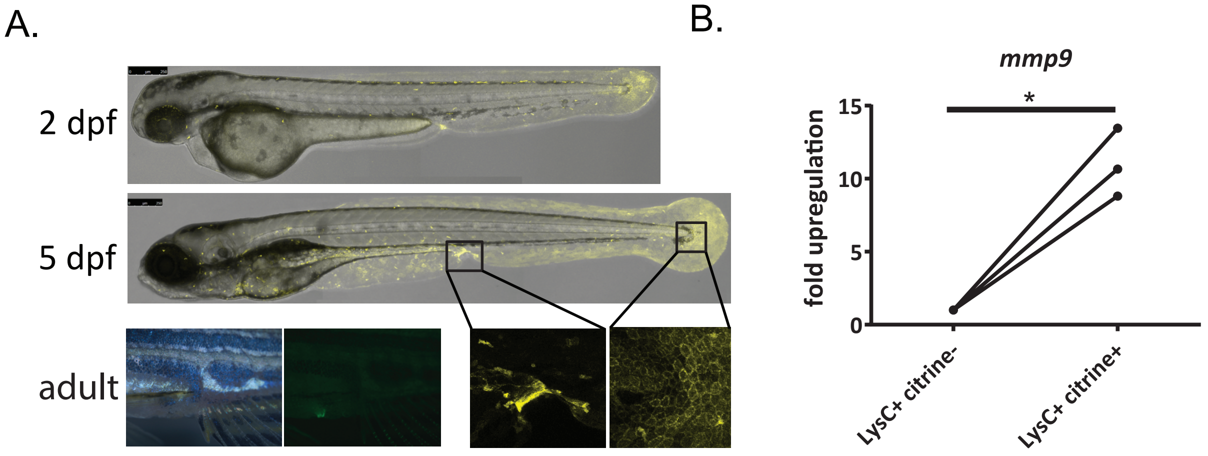

Supplementary figure 2:
Immune activation regulates frequency of Mmp9⁺ neutrophils

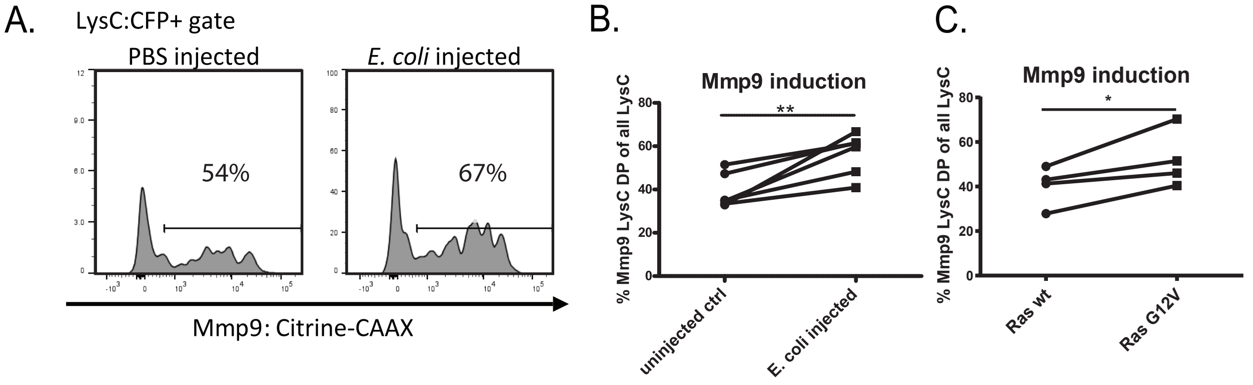

### Supplementary figure 3:
### Workflow and quality control of scRNA-seq of zebrafish kidney marrow neutrophils

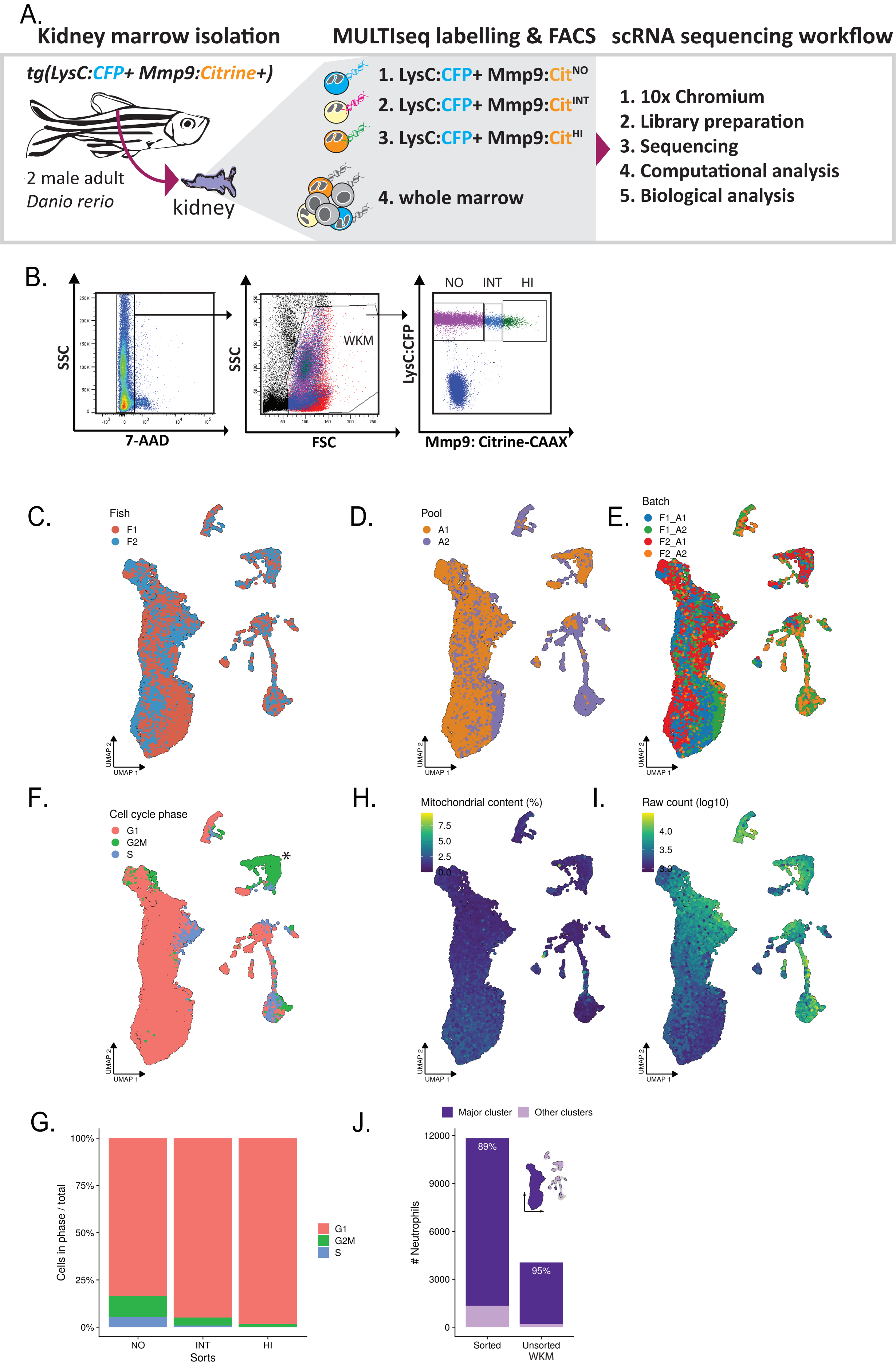

Supplementary Figure 4:
Identification of candidate regulators per gene module by dynamic time warping analysis

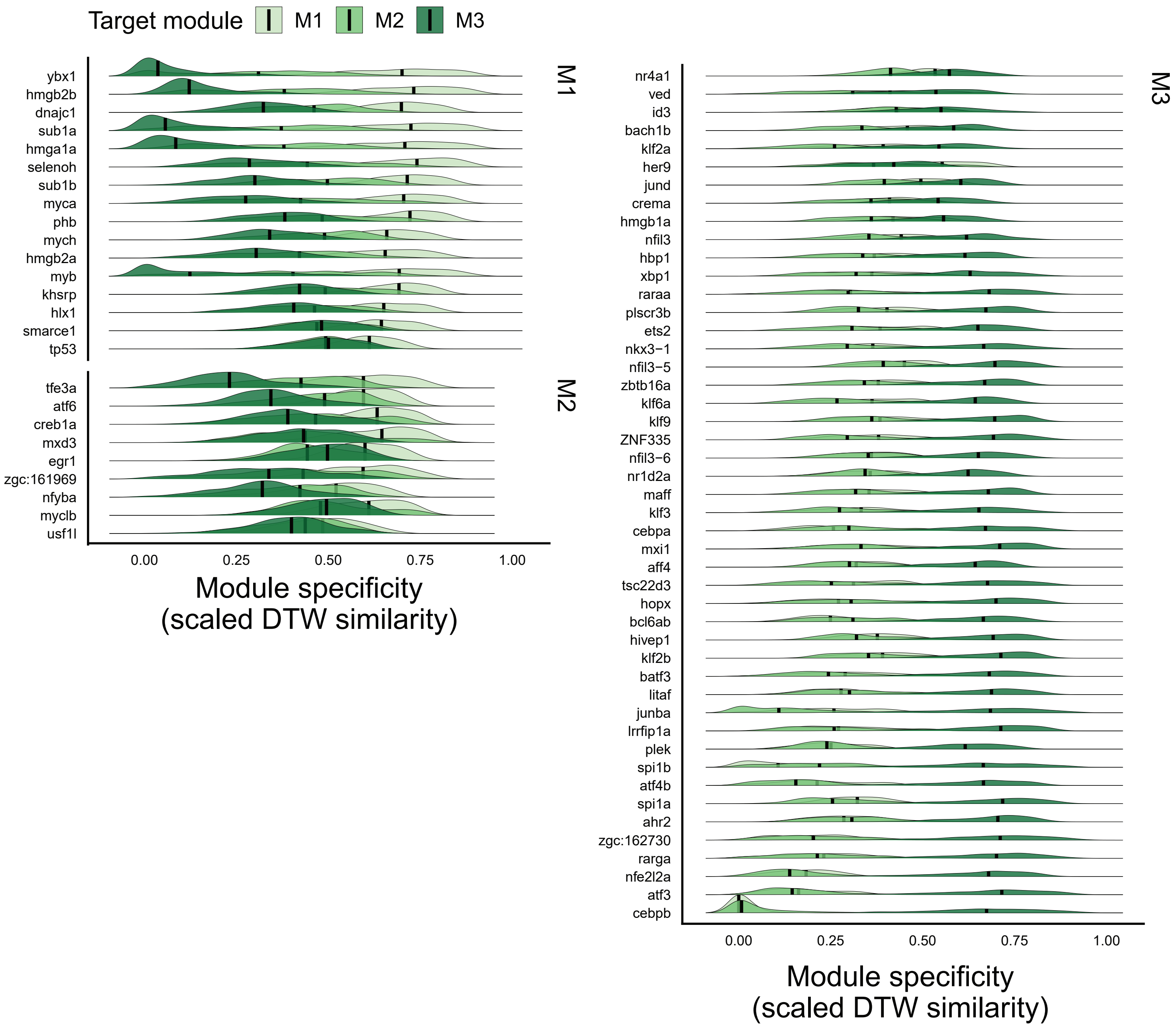

Supplementary Figure 5:
Cross-correlation analysis of gene expression during zebrafish and mammalian neutrophil maturation

A.Cross-correlation lags of selected genes relative to zebrafish

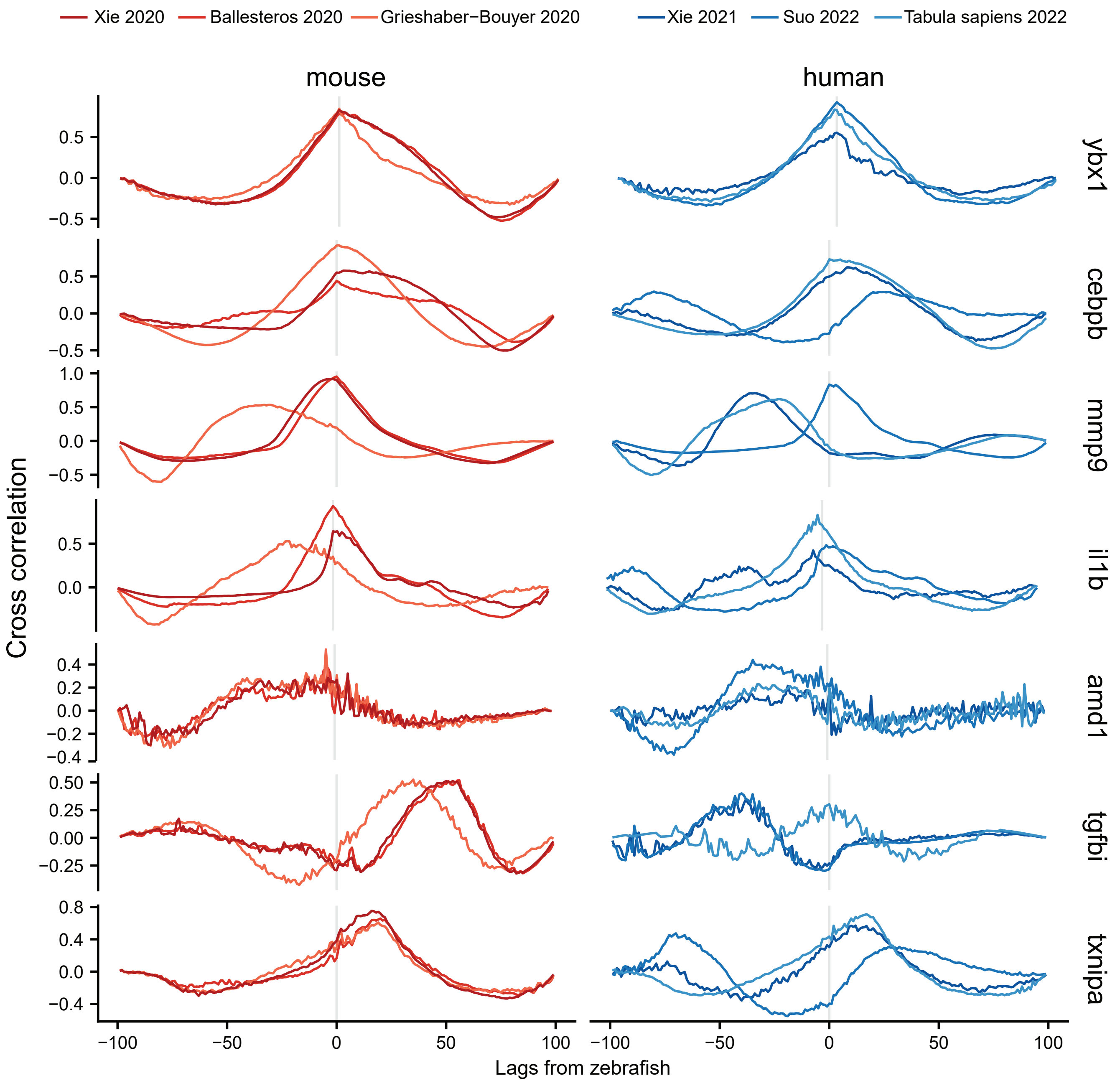

### Supplementary Figure 5:

#### B.Cross-correlation lags from zebrafish reference for all common differentially expressed genes (M1).

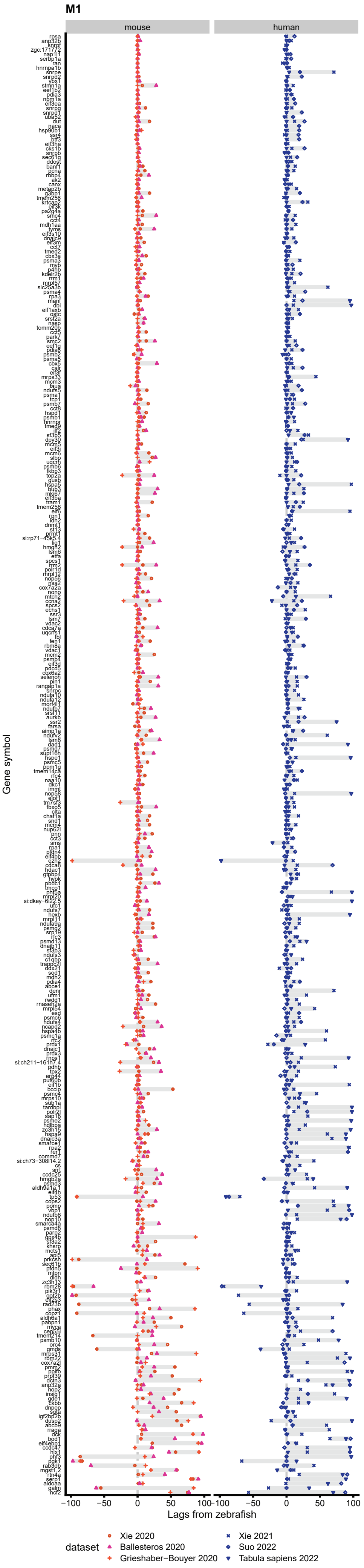

### Supplementary Figure 5:

#### B.Cross-correlation lags from zebrafish reference for all common differentially expressed genes (M2)

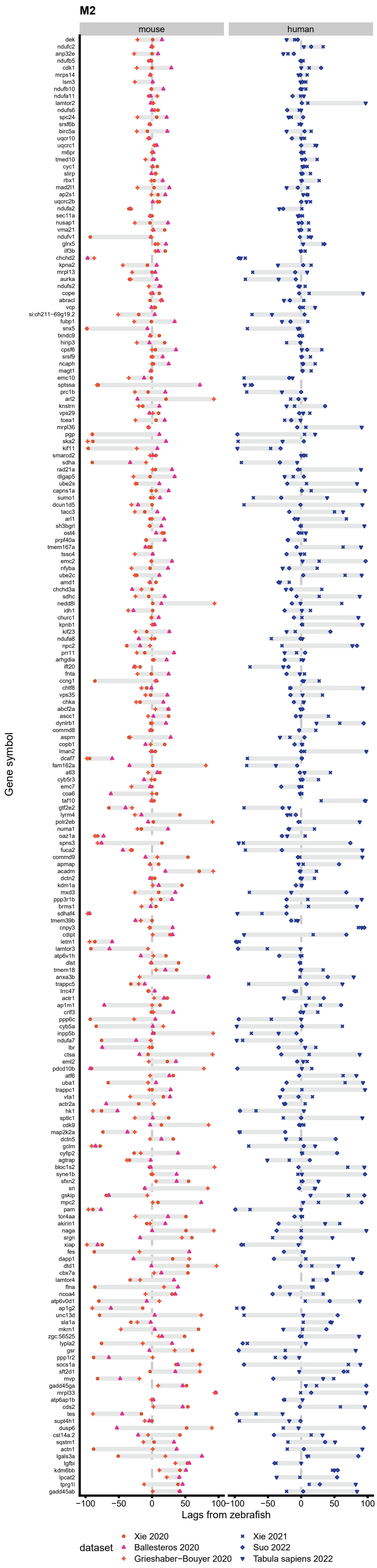

### Supplementary Figure 5:

#### B.Cross-correlation lags from zebrafish reference for all common differentially expressed genes (M3)

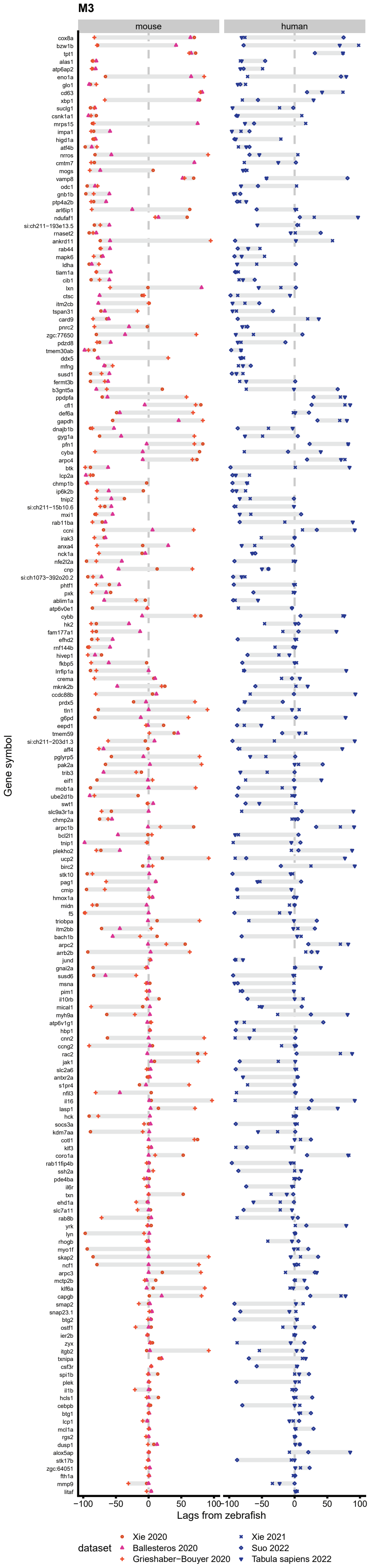

Supplementary Figure 6:

HAY_immature signature pre-dominantely includes late maturation genes

A. HAY signature genes included in the zebrafish signature

| HAY_zfish_common | zfish_module |
|---|---|
| ctnnb1 | $M1_{pan}$ |
| gpx4b | |
| ifi30 | |
| mhc1uba | |
| pfdn5 | |
| uba52 | |
| dusp6 | $M2_{pan}$ |
| lgals3a | |
| sec11a | |
| sgk1 | |
| atp6v1g1 | $M3_{pan}$ |
| b3gnt5a | |
| birc2 | |
| cebpb | |
| crip1 | |
| ctss2.1 | |
| hbegfa | |
| hk2 | |
| hmox1a | |
| hpse | |
| il1b | |
| irf2bp2b | |
| itm2ba | |
| lcp2a | |
| lgals2a | |
| mapk6 | |
| metrnla | |
| nfkbiaa | |
| notch2 | |
| ptp4a2b | |
| rhoab | |
| susd6 | |
| txnipa | |
| zgc:158343 | |

B. Application of HAY_immature signature on Ramirez et al. in-vitro differentiated neutrophils

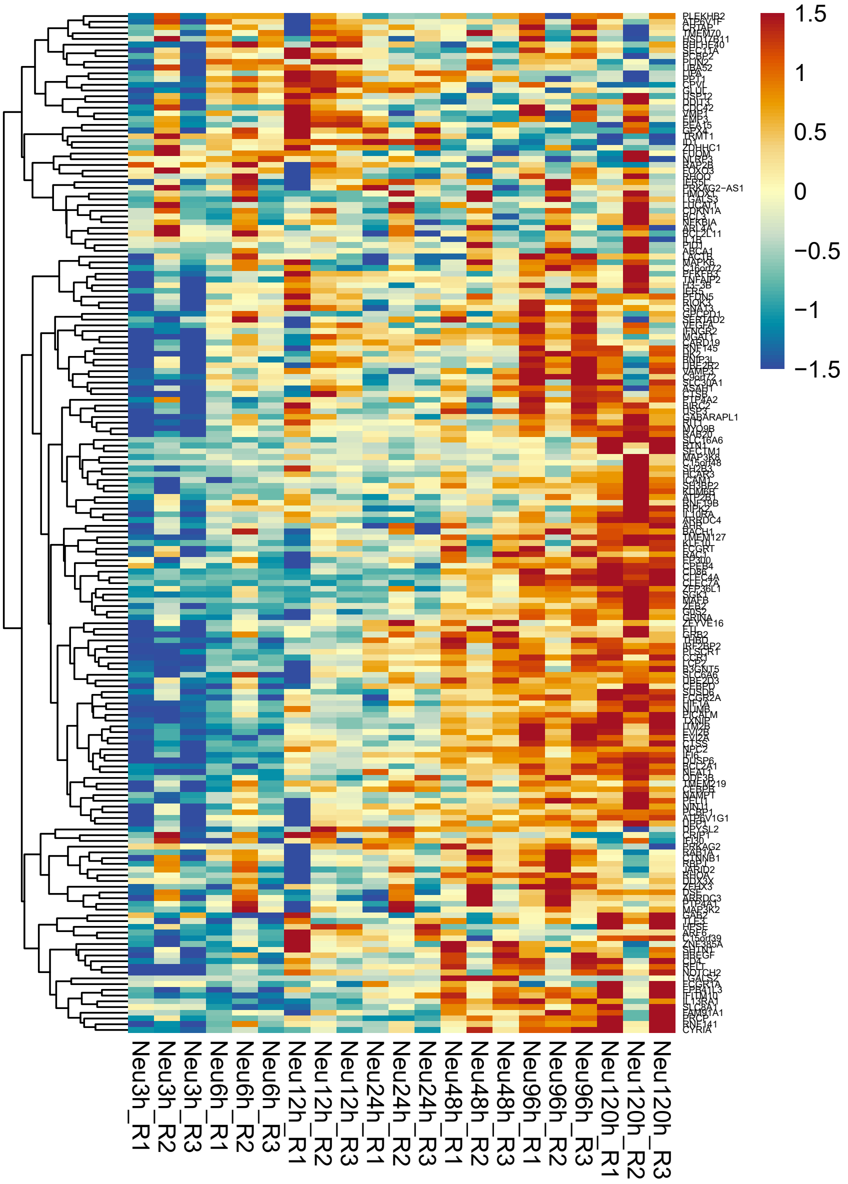

Supplementary Figure 7:
Human Neuroblastoma-BM RNA-seq analysis

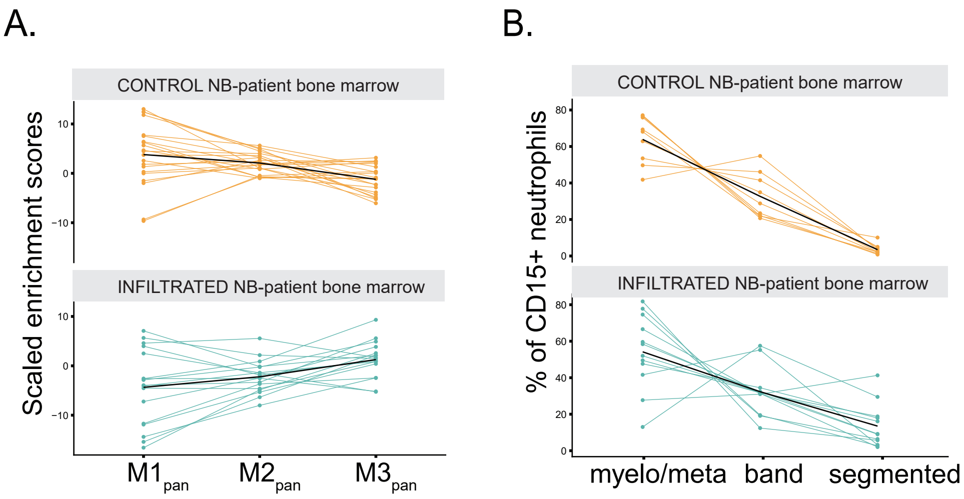

Supplementary Figure 8:
Optimal selection of the number of phases and modules

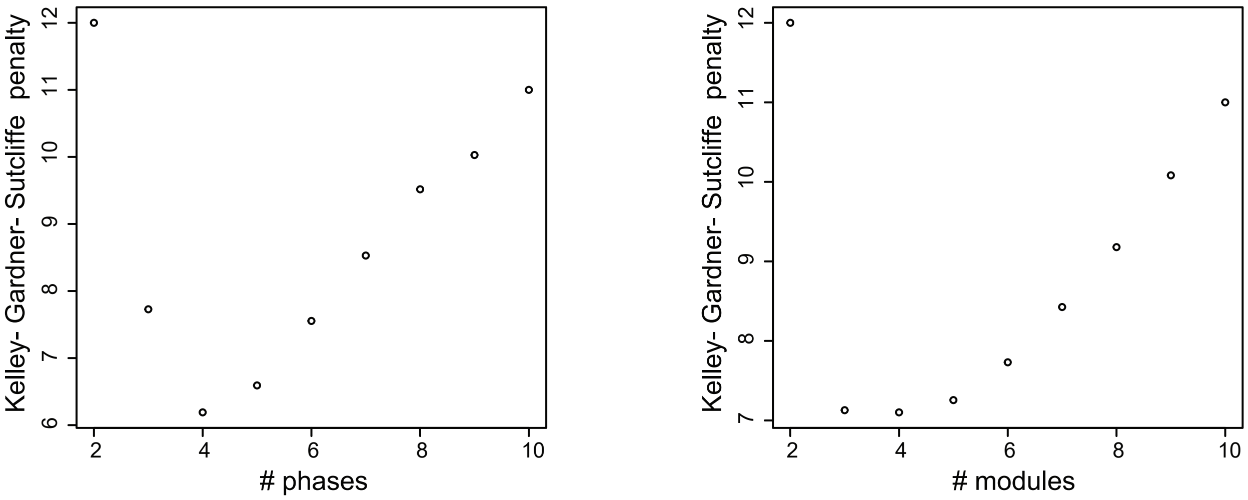

Supplementary Figure 9:

Model for neutrophil development in zebrafish based on scRNA sequencing.

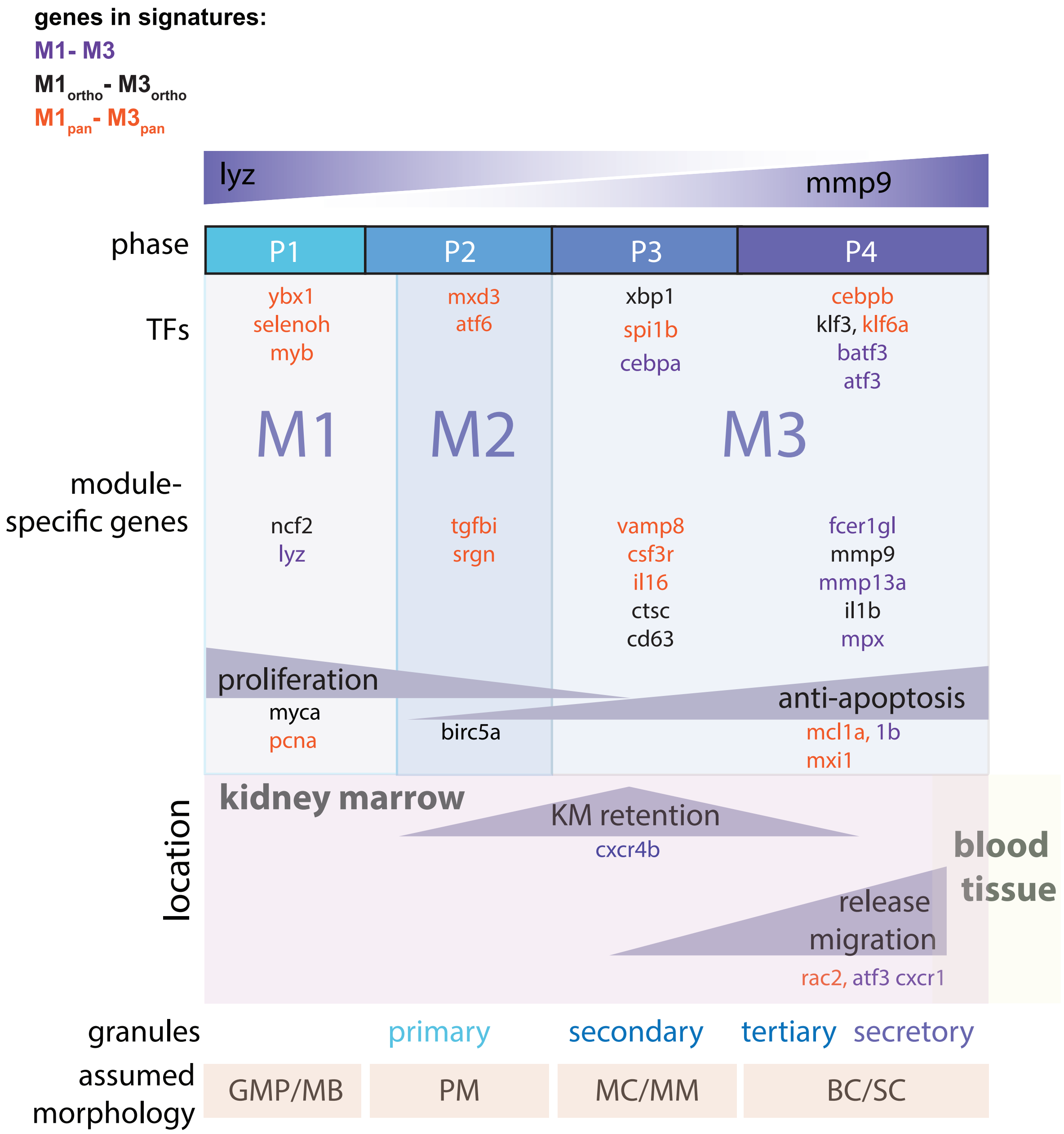
